# DWARF14 and KARRIKIN INSENSITIVE2 mediate signaling of the apocarotenoid zaxinone in Arabidopsis

**DOI:** 10.1101/2024.07.03.601835

**Authors:** Juan C. Moreno, Umar Shahul Hameed, Aparna Balakrishna, Abdugaffor Ablazov, Kit Xi Liew, Muhamad Jamil, Jianing Mi, Kawthar Alashoor, Alexandre de Saint Germain, Stefan T. Arold, Salim Al-Babili

## Abstract

The natural growth regulator zaxinone increases the levels of the phytohormones strigolactone (SL) and abscisic acid in Arabidopsis (*Arabidopsis thaliana*) via unknown mechanisms. We demonstrate that parts of the effects of zaxinone in Arabidopsis depend on the SL receptor DWARF14 (*At*D14), the karrikin receptor KARRIKIN INSENSITIVE2 (*At*KAI2), and the F-Box protein MORE AXILLARY BRANCHING2 (*At*MAX2) that mediates the signaling of SLs and karrikins. Binding assays and co-crystallization revealed zaxinone as an additional ligand of *At*D14 and an SL antagonist that interrupts the interaction of *At*D14 with *At*MAX2. Zaxinone also bound to *At*KAI2. These findings unveil a perception mechanism for zaxinone in Arabidopsis and demonstrate the capability of *At*D14 and *At*KAI2 to bind signaling molecules, other than strigolactones or karrikins, and mediate their transduction.

## Introduction

Strigolactones (SLs) are carotenoid-derived phytohormones with diverse roles in plant development and physiology (*1, 2*), including determining shoot and root architecture (*2, 3*). SLs are released by roots to allow the plant to communicate with symbiotic arbuscular mycorrhizal fungi (*4, 5*). These SLs also induce seed germination in root parasitic plants, such as *Striga hermonthica* (Striga), which precedes infestation by these weeds^2^.

SLs consist of a butenolide (D-ring) connected by an enol-ether bridge in *R*-configuration to a second, structurally variable moiety (*6*) represented by a tricyclic lactone ring (ABC-ring) in canonical SLs and different structures in non-canonical SLs. The structural diversity of SLs is linked to their specific biological roles (*7-9*). SLs are perceived by the α/β-fold hydrolase DWARF14 (D14), which binds to and cleaves them into the D-ring (which covalently attaches to the histidine residue of the D14 catalytic Ser-His-Asp triad) and the second moiety (*10, 11*). This attachment triggers the structural rearrangement of D14, allowing it to form a complex with the F-box protein MORE AXILLARY BRANCHING2/DWARF3 (MAX2/D3) and transcriptional repressors, such as SUPPRESOR OF MAX2-LIKE6 (SMXL6) in Arabidopsis (*Arabidopsis thaliana*) or D53 in rice (*Oryza sativa*). This process initiates the ubiquitination and degradation of the targeted transcriptional repressors by linking them to the MAX2-containing E3 ligase complex (*12*). D14 also perceives and hydrolyzes synthetic SL analogs and mimics, such as GR24, MP3, and Nijmegen-1 (*13, 14*).

A racemic mixture of the synthetic SL analog GR24, (±)-GR24, and (-)-GR24 bind to the D14 homolog KARRIKIN INSENSITIVE2 (KAI2) (*15*), which perceives karrikins (KARs). These small, non-hydrolysable butenolide smoke-derived compounds mimic an unidentified plant growth regulator termed KAI2-ligand (KL), which regulates seed germination, plant growth, biotic and abiotic stress responses, and mycorrhization (*16*). Interestingly, KAR signaling, which requires MAX2/D3 in Arabidopsis and rice (*16*), involves a perception mechanism similar to and overlapping with that of SLs (*17*). SUPRESSOR OF MAX2 1 (SMAX1) regulates seed germination and hypocotyl elongation via KAR signaling in Arabidopsis (*18, 19*), whereas SMAX1 regulates mesocotyl elongation in rice in the dark via its interaction with KAI2/D14L-D3 (*20*). These observations reflect the commonalities and specificities of the SL and KAR pathways in regulating different aspects of plant physiology.

The cleavage of carotenoids by reactive oxygen species or CAROTENOID CLEAVAGE DIOXYGENASES (CCDs) (*1, 21*) gives rise to apocarotenoid signaling molecules, e.g., β-cyclocitral, β-cyclocitric acid, β-ionone, anchorene, and zaxinone, which regulate plant growth, phytohormone homeostasis, metabolism, and stress responses (*22-26*). The effects of these compounds vary depending on the plant species. For instance, zaxinone treatment in rice resulted in reduced SL content, enhanced sugar uptake and metabolism, and improved root growth (*25, 27*), whereas in Arabidopsis, this treatment led to increased expression of the SL biosynthetic genes *CCD7* and *CCD8*, enhanced SL and abscisic acid (ABA) levels, and reduced root growth (*28*). Despite our understanding of the physiological effects of zaxinone and other apocarotenoids, their perception mechanisms remain elusive.

In this study, we demonstrated that zaxinone regulates the transcription of a wide range of genes in Arabidopsis, which partially depends on the receptors *At*D14 and *At*KAI2 as well as *At*MAX2-mediated signal transduction. Furthermore, the effect of zaxinone on SL biosynthesis requires *At*D14 and *At*MAX2. Zaxinone binds to both receptors and acts as an SL antagonist by competing for the binding cavity of *At*D14 and interrupting its interaction with *At*MAX2. Therefore, our findings identify a receptor for the apocarotenoid zaxinone and unveil the ability of the phytohormone receptors *At*D14 and *At*KAI2 to bind structurally and functionally different regulatory metabolites and transduce their signals.

## Results and Discussion

### *At*D14 is required for the responses of Arabidopsis to zaxinone

We previously showed that zaxinone treatment reduces SL contents by decreasing *CCD7* and *CCD8* transcription and enhanced root growth in rice cv. Nipponbare (*25*), which requires functional SL biosynthesis and perception^36^. To determine whether the effect of zaxinone on SL biosynthesis in Arabidopsis also relies on SL perception, we treated the SL receptor mutant *Atd14* with zaxinone and quantified the SL methyl carlactonoate (MeCLA) in root tissues using LC-MS. We also examined the activity of root exudates from zaxinone-treated plants in inducing seed germination in the root parasitic plant Striga (*28*). In contrast to wild-type (WT) Arabidopsis (*Col-0*), *Atd14* plants did not show increased MeCLA contents upon zaxinone treatment, indicating that this effect of zaxinone is dependent on *At*D14 (Fig. 1a). The results of the Striga bioassay were consistent with the results of LC-MS quantification (Fig. 1b-c; fig. S1a). We also tested root exudates of *Atkai2* and *Atmax2* plants following zaxinone treatment. Striga seed germination increased in response to treatment with exudates from *Atkai2* (and *Col-0*) but not *Atmax2* plants (Fig. S1b). To determine whether the effect of zaxinone on ABA content also depends on *At*D14, *At*MAX2, or *At*KAI2, we performed seed germination assays with the corresponding mutants in medium supplemented with or without zaxinone, or ABA as a positive control (Fig. 1d; fig. S2). Similar to ABA, zaxinone treatment delayed germination in all mutants, suggesting that neither *At*D14, *At*KAI2, nor *At*MAX2 is required for zaxinone-dependent increases in ABA content (Fig. 1d-e). ABA quantification in plants grown in hydroponic medium supplemented with zaxinone confirmed the finding that ABA accumulation in Arabidopsis is not dependent on *At*D14, *At*KAI2, or *At*MAX2 (Fig. 1f), pointing to the presence of other components for the perception and signal transduction of this apocarotenoid.

**Fig. 1.**
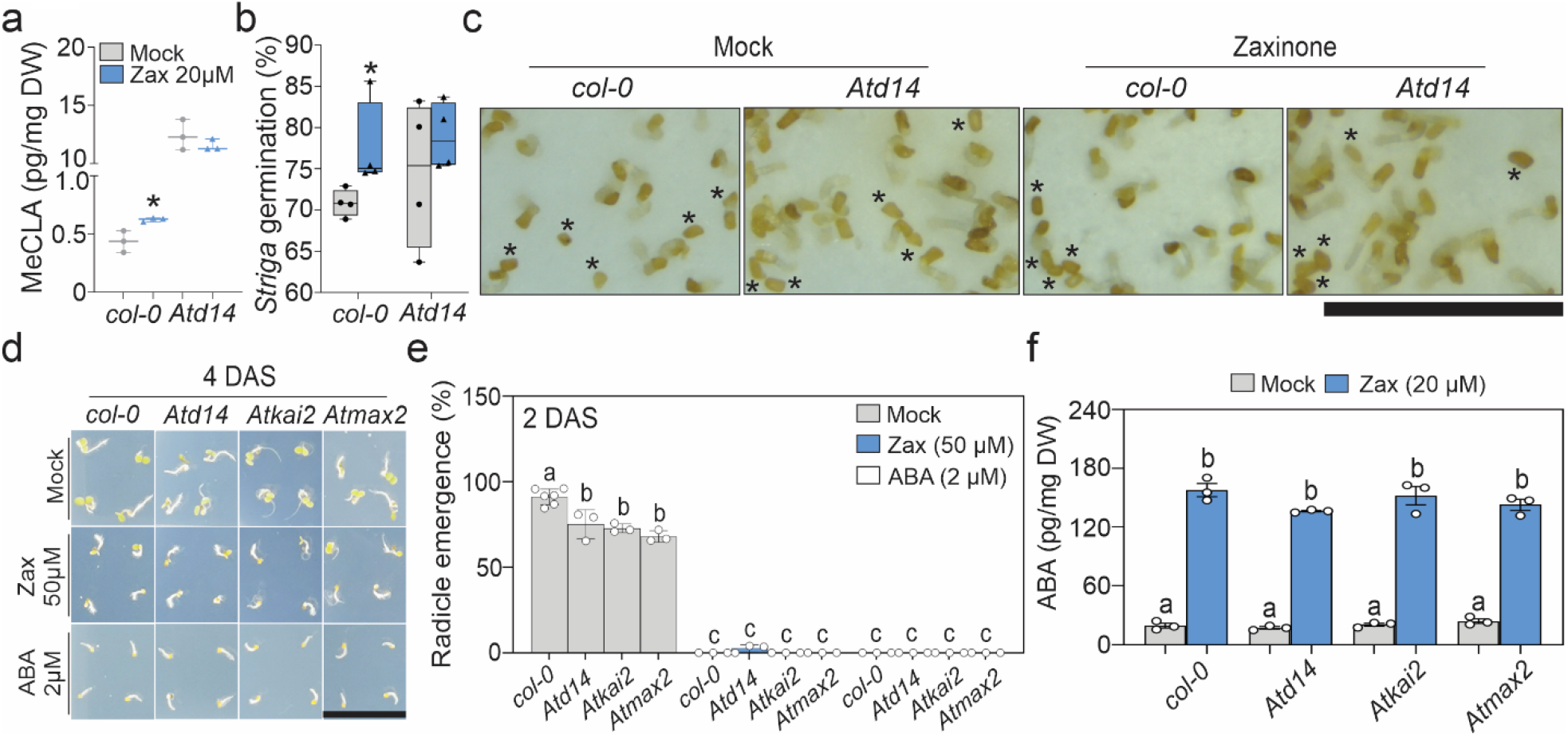
*At*D14 is required for zaxinone response in Arabidopsis. **a** Strigolactone (MeCLA) quantification in Arabidopsis roots subjected to Mock (acetone) or zaxinone treatment (20 *µ*M; *n*=3). **b-c** Striga germination bioassay using the root exudates from 5-weeks old WT *Col-0* and mutant *d14* Arabidopsis plants treated with mock (grey) or zaxinone (blue; 20 *µ*M) for 6 hours (*n*=4). In **c** photos are a zoom in of the discs where Striga seeds were germinated upon exudate application (scale bar=3 mm). For **a**-**b** unpaired two tails Student’s *t*-test were performed to assess significance (*: p<0.05). The complete discs can be found in Supplementary Fig. 1a. **d** Zoom in of 4 days-old Arabidopsis seedlings (DAS) germinated on mock, zaxinone and ABA-supplemented MS media. Scale bar: 3cm. **e** Quantification of seed germination for all genotypes and treatments two days after sowing (the experiment was repeated twice). Single data points represent measurements from three or six (for WT) independent plates with 30-35 seeds per genotype. **f** ABA quantification in the roots of 5-weeks old Arabidopsis *Col-0, Atd14, Atkai2*, and *Atmax2* plants treated with mock (1% DMSO) and zaxinone for six hours (*n*=3). In **e**-**f** letters denote significance assessed by one way ANOVA (*n*=3).

### D14 and MAX2 are required for the zaxinone-dependent induction of SL biosynthetic genes in Arabidopsis

We analyzed the root transcriptomes of *Col-0, Atd14, Atkai2*, and *Atmax2* plants (Fig. 2a) upon treatment with 20 **µ**M zaxinone via RNA sequencing (RNA-seq). We identified 1835 differentially expressed genes in *Col-0* (DEGs; Log_2_ F.C.>1 (up) or Log_2_ F.C.< -1 (down), padj<0.05; Data S1-2), which were enriched in a wide range of biological processes in different cellular compartments (fig. S3; Data S3-10; Fisher test and Bonferroni post-test, p>0.05). Consistent with our previous study (*28*), we observed significant increases in the transcript levels of *CCD7* (Log_2_ F.C.>2) and *CCD8* (Log_2_ F.C.>1.5), encoding proteins that mediate SL biosynthesis (fig. S3c; Data S1-2). We searched for genes whose transcription was affected by zaxinone in WT (1835 genes; fig. S3c; Data S2) but not in *Atd14, Atkai2*, or *Atmax2* plants (Data S11-16). By comparing *Col-0* zaxinone (Z) vs. mock (M) with *Atd14* Z vs. M, we identified 340 DEGs (among the 1835 zaxinone-responsive DEGs in *Col-0*) that did not respond to zaxinone treatment in *Atd14* (-1>Log_2_F.C./1<Log_2_ F.C. and padj>0.05; Fig. 2b; Data S17-19), suggesting the need for a functional *At*D14 receptor for the zaxinone-dependent regulation of these genes. Similarly, 458 and 486 genes required functional *At*KAI2 (-1>Log_2_F.C./1<Log_2_ F.C. and padj>0.05; Fig. 2b; Data S20-22) and *At*MAX2 (-1>Log_2_F.C./1<Log_2_ F.C. and padj>0.05; Fig. 2b; Supplementary Datasets 23-25) for their zaxinone-dependent regulation, respectively. Moreover, 205 genes were dependent on *At*D14, *At*KAI2, and *At*MAX2 for their zaxinone-dependent regulation (fig. S4), whereas 56, 103, and 40 genes required the pairs D14-MAX2 (Data S26-27), KAI2-MAX2, and D14-KAI2, respectively, for their zaxinone-dependent regulation (fig. S4). Additionally, 39, 110, and 122 genes were solely dependent on *At*D14, *At*KAI2, or *At*MAX2, respectively, for their zaxinone-dependent regulation (fig. S4), suggesting that these proteins also function independently in different signaling networks. We validated the responses of four of the 39 genes that were dependent only on *At*D14 for their zaxinone-dependent regulation (Data S28-29) using reverse transcription quantitative PCR (RT-qPCR) (fig. S5). Among these genes, ABA RESPONSIVE (*ABR*) expression was induced by zaxinone treatment in *Col-0, Atkai2*, and *Atmax2* but not in *Atd14* confirming the possibility that the transcription of these set of genes only require D14 and perhaps an unidentified ubiquitin E3 ligase different to MAX2. This is in line with the finding showing that the rice ortholog *Os*D14 interacts with a RING-finger ubiquitin E3 ligase (SDEL1) under phosphate deficiency, leading to the degradation of SPX DOMAIN-CONTAINING PROTEIN 4 and the release of PHOSPHATE STARVATION RESPONSE PROTEIN 2 (*29*).

**Fig. 2.**
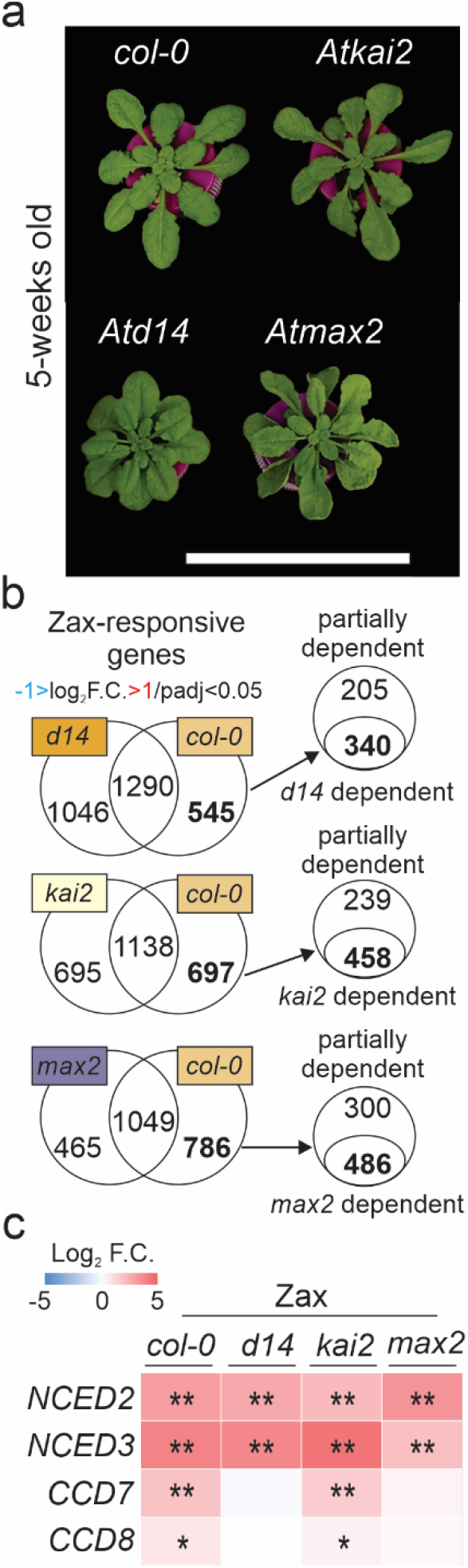
RNAseq analysis of Arabidopsis *Col-0, d14, kai2* and *max2* roots upon zaxinone treatment. **a** Plant phenotypes of 5-weeks old Arabidopsis *Col-0* and mutants grown in hydroponic media. **b** Venn diagrams showing zaxinone (Zax) responsive genes (DEGs) measured by RNAseq in *Atd14, Atkai2*, and *Atmax2* mutant backgrounds in response to zaxinone (20 *µ*M). The expression of 545, 697 and 786 DEGs (up and down) was modulated by zaxinone in the wild type, however the expression of these DEGs remained unchanged in the *Atd14, Atkai2* and *Atmax2* mutant backgrounds, respectively. **c** Heatmap representation showing the expression patterns of genes involved in ABA and SL biosynthesis in *Col-0, Atd14, Atkai2* and *Atmax2* mutant backgrounds. All selected DEGs fulfill the threshold Log_2_ F.C.>1 (up) or Log_2_ F.C.< -1 (down) and padj<0.05. The RNAseq experiment was performed by applying 20 *µ*M zaxinone to the hydroponic media of 5-weeks old Arabidopsis plants for 6 hours (*n*=4).

Consistent with the results of *in vivo* tests, LC-MS analysis, and the Striga bioassay (Fig. 1), zaxinone upregulated the ABA biosynthetic genes *9-CIS-EPOXYCAROTENOID DIOXYGENASE* (*NCED*) *AtNCED2* and *AtNCED3* (padj<0.05) in all lines examined, i.e., *Col-0, Atd14, Atkai2*, and *max2* (Fig. 2c). However, the SL biosynthetic genes *AtCCD7* and *AtCCD8* were upregulated only in *Col-0* and *Atkai2* but not in *Atd14* or *Atmax2* (Fig. 2c). RT-qPCR further confirmed the responses of *AtCCD7, AtCCD8*, and *AtNCED3* in *Col-0, Atd14, Atkai2*, and *Atmax2* (fig. S6). These results confirm that the effect of zaxinone on ABA biosynthesis is independent of SL and KAR perception, whereas its effect on SL biosynthesis requires a functional *At*D14 and *At*MAX2 but not *At*KAI2.

### *In vitro* zaxinone-*At*D14 interaction and co-crystallization reveal zaxinone as an SL-antagonist

We examined whether zaxinone directly binds to *At*D14 and *At*KAI2 (Fig. 3) by performing nano differential scanning fluorescence (nanoDSF) experiments with purified *At*D14 and *At*KAI2. Zaxinone treatment increased the melting temperature (*T*_*m*_) of *At*D14 by 3±0.23 to 5.2±0.11 °C, whereas (±)-GR24 treatment decreased the *T*_*m*_ of *At*D14 by 7±0.11 to 7.9±0.10 °C, both at concentrations of 10 *µ*M and above (Fig. 3a and fig. S7a; Table S1-3). Zaxinone also stabilized *At*KAI2 but only at concentrations of 50 and 100 *µ*M (fig. S7b; Table S4-5). We also tested the *At*D14 and *At*KAI2 homolog *At*D14-LIKE2 but failed to detect any effect of zaxinone (fig. S7c and Table S6-7). Incubation with the karrikin KAR2 did not affect the *T*_*m*_ of *At*KAI2 (fig. S7d; Table S1), which is in line with previous studies (*30, 31*).

**Fig. 3.**
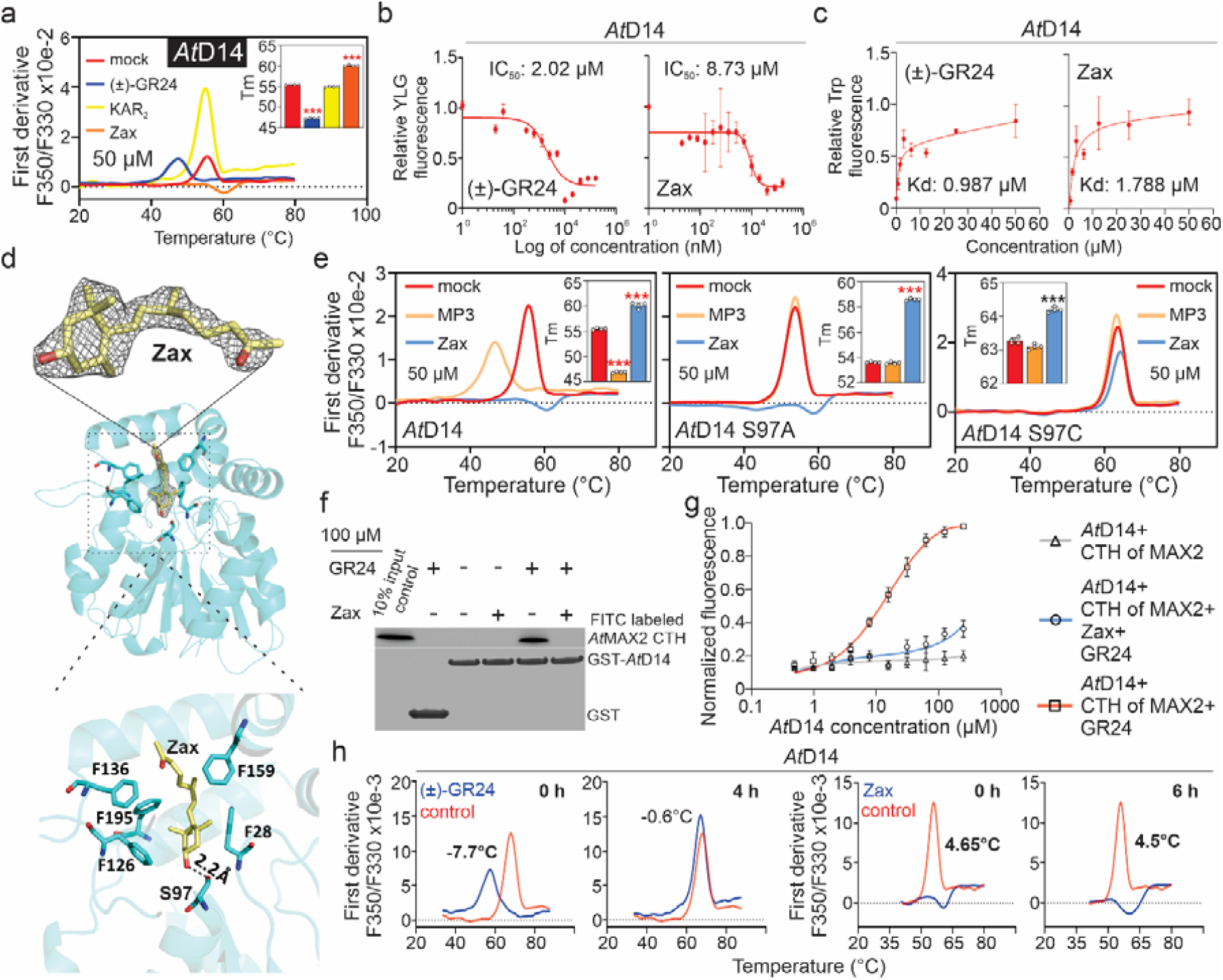
*In vitro* and co-crystallization experiments identify zaxinone as an *At*D14 ligand. **a** Nano differential scanning fluorimetry (nanoDSF) assays for *At*D14 (6 *µ*M) in the absence and presence of *rac*-GR24, KAR_2_, and zaxinone. **b** YLG cleavage by *At*D14 (3 *µ*M) in the presence of increasing (±)-GR24 and zaxinone concentrations. Data are the mean ± SE, *n=3*. **c** Binding properties of *At*D14 (10 *µ*M) in the presence of (±)-GR24 and zaxinone. Apparent dissociation constant (*K*_*d*_) were derived from intrinsic protein fluorescence measurements at increasing substrate concentrations. Data are the mean and error ± SD (*n=3*). **d** X-ray crystallographic structure of the *At*D14-zaxinone complex. Top left: ribbon overview of *At*D14 (cyan) with zaxinone (stick representation, carbons in yellow and oxygen in red) bound in its active site. Gray mesh shows the 2.4 Å electron density 2Fo-Fc omit map, contoured at 1 σ. Six key amino acid residues for zaxinone binding are shown as sticks (F28, S97, F126, F136, F159 and F195). Bottom left: zoomed view of the zaxinone model (yellow) fitted into the 2Fo-Fc electron density omit map. Right: zoomed view into the zaxinone binding site. Dotted line illustrates the 2.2 Å hydrogen bond from the active site S97 to the OH group of zaxinone. **e** NanoDSF specificity assay for *At*D14, *At*D14 S97A and *At*D14 S97C (all proteins ∼6 *µ*M) in the absence and presence of the SL analog MP3 and zaxinone (50 *µ*M). **f** Pull-down of the FITC labeled CTH peptide of *At*MAX2 with GST-*At*D14 in the absence and presence of (±)-GR24and/or zaxinone. **g** Competition assay between *At*D14 and CTH peptide of *At*MAX2 in the absence and presence of (±)-GR24 and/or zaxinone. **h** Evaluation of *At*D14 substrate stability. Melting temperature curves of *At*D14 (6 *µ*M) with or without pre-incubation with (±)-GR24 (50 *µ*M) and zaxinone (20 *µ*M) for the indicated time period. Tm values (a, e, i) were calculated using default settings in PrometheusNT.48 software (means ± SE, *n=4*). *** p<0.0005 (unpaired two tails Student *t*-test). P-value (<0.05) and a shift in the melting profile of at least 1°C were used to define binding (red asterisks).

We then investigated whether zaxinone is a competitor of SL in binding to *At*D14 using a Yoshimulactone (YLG) hydrolysis assay; (±)-GR24 served as a positive control. The half-maximal inhibitory concentration (IC_50_) of (±)-GR24 was 2.0 *µ*M, while that of zaxinone was 8.7 *µ*M, indicating that zaxinone is a competitor of SLs, although with a lower efficiency than the SL analog (±)-GR24 (Fig. 3b). We also determined the dissociation constants (*K*_*d*_) for both compounds using an intrinsic tryptophan fluorescence assay (Fig. 3c). *At*D14 bound to (±)-GR24 and zaxinone with a *K*_*d*_: 0.99 *µ*M and *K*_*d*_: 1.8 *µ*M, respectively (Fig. 3c).

Next, we crystalized *At*D14 in the presence of zaxinone. These crystals diffracted X-rays to determine the ⏢ 2.4 resolution. The crystal structure was determined by molecular replacement using the *At*D14 structure as a template (PDB accession code 4IH4; Fig. 3d, Table S8). Clear electron density was visible in the SL binding pocket of *At*D14, into which the zaxinone molecule could be unambiguously fitted (Fig. 3d). Six amino acids were crucial for zaxinone binding: F136, F195, F126, S97, F28, and F159. The OH group of zaxinone was present at a distance of 2.2 ⏢ from the catalytic triad residue S97, which indicates that the interaction may be stabilized via the formation of a hydrogen bond (Fig. 3d). To assess the importance of this hydrogen bond, we tested the effect of zaxinone on S97A and S97C mutants of *At*D14 using a nanoDSF assay; the SL analog methyl phenlactonoate 3 (MP3) (*32*) served as a positive control (Fig. 3e). Similar to (±)-GR24, MP3 destabilized wild-type *At*D14 by 8.6±0.11 °C, whereas zaxinone stabilized it by 4.6±0.29 °C (Fig. 3e). The *At*D14 S97A mutation impeded the interaction between MP3 and *At*D14 (Fig. 3e), in agreement with previous findings for GR24 (*11*), but did not profoundly affect the interaction with zaxinone (*T*_*m*_: 5±0.05 °C). By contrast, the S97C mutation of *At*D14 severely decreased its interactions with both MP3 (0.18±0.06 °C shift in *T*_*m*_) and zaxinone (0.9±0.06 °C shift in *T*_*m*_; Fig. 3e). According to the crystal structure, the S97C mutation would result in an unfavorable proximity of the zaxinone OH moiety with the hydrophobic sulfur of the cysteine in addition to shortening the distance between the side chain of position 97 and the hydroxyl group of zaxinone (fig. S8). This analysis further corroborated the crystallographic model for the association of zaxinone with the *At*D14 active site.

Considering that the hydrolysis of GR24 is associated with a conformational change in D14 that triggers MAX2 binding (*12*), we asked whether zaxinone antagonizes SL signaling by preventing *At*D14 from binding to *At*MAX2. To investigate this possibility, we performed a pull-down assay to examine the ability of *At*D14 to associate with a C-terminal helix (CTH) of *At*MAX2 upon SL binding (*12*). GST-D14 bound to the fluorescein isothiocyanate (FITC)-labeled *At*MAX2 CTH peptide in the presence of (±)-GR24. However, this interaction did not take place when *At*D14 was incubated with zaxinone prior to adding (±)-GR24 (Fig. 3f). To support these findings, we performed a competition assay in which we measured the fluorescence polarization of FITC-CTH in the presence of increasing *At*D14 concentrations as an indicator of CTH-*At*D14 complex formation (Fig. 3g). *At*D14 did not interact with CTH in the absence of (±)-GR24 (gray line; Fig. 3g). Adding (±)-GR24 resulted in a strong increase in FITC-CTH fluorescence polarization, which is indicative of binding (red line; Fig. 3g). However, incubation of *At*D14 with zaxinone before adding (±)-GR24 almost completely blocked the CTH-*At*D14 association, as revealed by the lack of increase in fluorescence polarization. Collectively, these results demonstrate that zaxinone interferes with the downstream signaling of *At*D14 by competitively occupying its SL binding site without introducing SL-associated conformational changes. Interestingly, zaxinone has opposite effects on SL biosynthesis in Arabidopsis and rice (*25, 28*), although functional SL perception is required for its effects in both species (*25*).

Similar to *At*D14, zaxinone treatment increased the *T*_*m*_ of *Os*D14 at different *µ*M concentrations, whereas MP3 treatment led to a decrease in *T*_*m*_ (fig. S9a). Moreover, tryptophan fluorescence assays confirmed the direct interaction between *Os*D14 and zaxinone (fig. S9b). Additionally, the binding between *Os*D14 and zaxinone remained stable for up to two hours, whereas the degradation of MP3 was detected after just one hour (fig. S9c). However, our YLG hydrolysis assay did not indicate any competition between zaxinone and (±)-GR24 (fig. S9d) for binding with *Os*D14, suggesting disparities in the molecular interactions between *Os*D14-zaxinone and *At*D14-zaxinone. This observation is consistent with the differing responses of rice and Arabidopsis to zaxinone treatment.

If zaxinone cannot be cleaved, its effect on *At*D14 should last longer than that of (±)-GR24, which is hydrolyzed and released. To experimentally confirm this notion, we performed a time-course nanoDSF assay in which we incubated *At*D14 with 50 *µ*M (±)-GR24 for 0, 0.5, 1, 2, and 4 h or with 20 *µ*M zaxinone for 0, 1, 2, 4, and 6 h. We observed a clear interaction between (±)-GR24 (destabilization by 7.7±0.11 °C) and zaxinone (stabilization by 4.7±0.05 °C) with *At*D14 at 0 h. The (±)-GR24-associated *T*_*m*_ rapidly decreased from 7.7±0.11 °C to 1.15±0.05 °C within 2 h and dropped to 0.6±0.05 °C at 4 h, indicating that the hydrolyzed (±)-GR24 was released (Fig. 3h; fig. S10a and Table S9). By contrast, the *T*_*m*_ shift caused by zaxinone treatment remained unaltered throughout the measurement period (above 4 °C; Fig. 3h; fig. S10a and Table S10). Similar results were obtained when we examined the interaction of zaxinone with *At*KAI2 over time (fig. S10b and Tables S11-12). Moreover, *in silico* docking placed zaxinone in the active site of *At*KAI2 with a binding pose matching the crystallographic *At*D14-zaxinone complex, suggesting that zaxinone binds to and affects *At*KAI2 and *At*D14 in a similar manner (fig. S11a-c). Indeed, zaxinone treatment repressed the expression of several *At*KAI2-dependent genes, including *DLK2, SMXL2*, and *KUF1*, in *Col-0* but not *Atkai2* plants (Data S2, 14).

Altogether, our transcript data reveal a KAI2-specific response to zaxinone (Data S30-32), and our *in vitro* and *in silico* data suggest that zaxinone could be a candidate for a *At*KAI2 ligand that acts as an antagonist of the sought-after KL, indicating the ability of KAI2 to perceive different endogenous signals. Notably, the volatile sesquiterpenoid (-)-germacrene D, which lacks a butenolide ring, was recently shown to bind to KAI2ia (an intermediate KAI2 clade receptor) and trigger a signaling cascade that regulates plant fitness in petunia (*Petunia hybrida*) (*33*). In *Medicago truncatula* and barley (*Hordeum vulgare*), Nodulation Signaling Pathway (NSP) transcription factors likely regulate the production of small molecules required for activating *Os*D14L (KAI2) and the subsequent suppression of SMAX1 (*34*). Surprisingly, 77% of the genes regulated by NSPs in *M. truncatula* identified in a recent study were involved in SL, carotenoid, and apocarotenoid biosynthesis (e.g., the zaxinone synthase gene), suggesting that KL might have its origin in one of these pathways (*34*).

SL biosynthesis is governed by a negative feedback mechanism triggered by SLs and analogs, which downregulate the transcription of SL biosynthetic genes (*35*). This mechanism is supported by the finding that disrupting the SL receptor D14 or the SL signaling component MAX2/D3 increased the levels of SLs and related biosynthetic transcripts (*25, 36*). Taken together, our results allowed us to uncover a mechanism in which the apocarotenoid zaxinone binds to the binding pocket of the SL receptor *At*D14, increasing the expression of SL biosynthetic genes (*CCD7* and *CCD8*), and ultimately SL contents, by interfering with the negative feedback loop that requires the binding of SLs to *At*D14. This hypothesis explains why *At*MAX2 is required for the effect of zaxinone on SL biosynthesis and the relationship between SL and zaxinone responses.

## Conclusion

In Arabidopsis, the effects of the apocarotenoid zaxinone are largely mediated by the SL receptor *At*D14, the KAR receptor *At*KAI2, and their downstream signaling component *At*MAX2. Despite its structural difference from SLs, zaxinone competitively binds to the active site of *At*D14. Thus, zaxinone acts as a long-lasting, non-hydrolyzable antagonist that opposes downstream signaling mediated by *At*D14 in the presence of SLs by blocking the interaction between *At*D14 and *At*MAX2. Taken together, our results reveal a receptor for zaxinone and provide evidence that the butenolide hormone receptors *At*D14 and *At*KAI2 channel different endogenous signals in Arabidopsis. This finding opens a new avenue in plant hormone research and might explain why the sought-after endogenous ligand with karrikin activity is still elusive.

## Supporting information

All supplemental material including methods, Figures and Tables

## Acknowledgements

We are thankful to Lamis Berqdar, Vijayalakshmi Ponnakanti, Gadah Alamri, and Alice Stra for technical assistance in some of the plant-related work. We are grateful with Dr. Yoshiya Seto for providing the expression vectors carrying Arabidopsis D14 with single point mutations and Dr. Tom Bennett for providing *Col-0, Atd14-1, Atkai2-2*, and *Atmax2-1* seeds. We acknowledge SOLEIL for provision of synchrotron radiation facilities under proposal ID 20210195 and we would like to thank P. Montaville for assistance in using beamline PROXIMA 1. Research reported in this publication was supported by baseline funding, the Competitive Research Grants CRG2020 and CRG2022, and the grant REP/1/3842-01-01 given to S. Al-B from King Abdullah University of Science and Technology (KAUST).

## Author contributions

S. Al-B., J.C.M., U.F.S.H., and S.T.A. conceived and designed research. J.C.M., A.B., and A.A. performed zaxinone and GR24 treatments for RNAseq and hormone quantification experiments. J.C.M. and A.B. performed RNA extraction for RNAseq and qPCR experiments. J.C.M. analyzed and interpreted RNAseq results. A.A. and J.C.M. performed cDNA synthesis. A.A. and J.C.M. performed qPCR experiments. A.B. generated final expression vectors for protein expression. J.C.M., U.F.S.H., and A.B. performed protein expression and purification. J.C.M. performed nanoDSF, YLG, and tryptophan (trp) fluorescence assays with the assistance of A.B., K.A. performed YLG and trp assays for rice *Os*D14, and U.F.S.H. performed the pull-down and competition assay. U.F.S.H. and S.T.A. performed X-ray crystallography analysis, interpretation, and deposition. U.F.S.H performed *in silico* and guided modelling of interactions. J.M. and K.X.L. performed hormone quantification. M.J. performed the collection of exudates and *Striga* bioassays. J.C.M. performed seed germination assays. A.de S.G. generated the preliminary nanoDSF results indicating *At*D14-zaxinone binding. J.C.M. wrote the paper with input from M.J. and U.F.S.H. S.T.A and S. Al-B. edited the paper and supervised the project.

## Competing interests

Authors declare that they have no competing interests.

## Data availability

All RNAseq data were deposited in OMNIBUS. Sequence data from this article can be found on TAIR under the following accession numbers: *AtMAX3*/*CCD7* (AT2G44990), *AtMAX4*/*CCD8* (AT4G32810), *AtMAX2* (AT2G42620), *AtNCED3* (AT3G14440), *AtD14* (AT3G03990), *AtKAI2* (AT4G37470), *AtGIK* (AT2G35270), *AtABR* (AT3G02480), *AtSnRK2-7* (AT4G40010), and *AtNPF3*.*1* (AT1G68570). Co-ordinates for the crystal structure was deposited in PDB with the accession code 8I7Y. All analysis and results are available in the Supplementary Information and Datasets (datasets are available in the following link:https://docs.google.com/spreadsheets/d/1CIngfYpmTpuSxDKt8LoLFxw817AbBQLR/edit?usp= sharing&ouid=105616710644855832961&rtpof=true&sd=true).

## Supplementary Materials

Materials and Methods

Figs. S1 to S11

Tables S1 to S13

References (28, *37*–56)

Data S1 to S32

(https://docs.google.com/spreadsheets/d/1CIngfYpmTpuSxDKt8LoLFxw817AbBQLR/edit?usp=sharing&ouid=105616710644855832961&rtpof=true&sd=true)

## Notes

### Competing Interest Statement

The authors have declared no competing interest.

## References

1. J. C. Moreno, J. Mi, Y. Alagoz, S. Al-Babili, Plant apocarotenoids: from retrograde signaling to interspecific communication. Plant J 105, 351–375 (2021).

2. S. Al-Babili, H. J. Bouwmeester, Strigolactones, a novel carotenoid-derived plant hormone. Annu Rev Plant Biol 66, 161–186 (2015).

3. M. T. Waters, C. Gutjahr, T. Bennett, D. C. Nelson, Strigolactone Signaling and Evolution. Annual Review of Plant Biology, Vol 68 68, 291–322 (2017).

4. K. Akiyama, K. Matsuzaki, H. Hayashi, Plant sesquiterpenes induce hyphal branching in arbuscular mycorrhizal fungi. Nature 435, 824–827 (2005).

5. C. Gutjahr, Phytohormone signaling in arbuscular mycorhiza development. Current Opinion in Plant Biology 20, 26–34 (2014).

6. A. de Saint Germain et al., An histidine covalent receptor and butenolide complex mediates strigolactone perception. Nature Chemical Biology 12, 787-+ (2016).

7. S. Ito et al., Canonical strigolactones are not the major determinant of tillering but important rhizospheric signals in rice. Sci Adv 8, eadd1278 (2022).

8. J. Y. Wang, G. E. Chen, J. Braguy, S. Al-Babili, Distinguishing the functions of canonical strigolactones as rhizospheric signals. Trends Plant Sci, (2024).

9. G. E. Chen et al., Disruption of the rice 4-DEOXYOROBANCHOL HYDROXYLASE unravels specific functions of canonical strigolactones. Proc Natl Acad Sci U S A 120, e2306263120 (2023).

10. C. Hamiaux et al., DAD2 is an alpha/beta hydrolase likely to be involved in the perception of the plant branching hormone, strigolactone. Curr Biol 22, 2032–2036 (2012).

11. Y. Seto et al., Strigolactone perception and deactivation by a hydrolase receptor DWARF14. Nat Commun 10, 191 (2019).

12. N. Shabek et al., Structural plasticity of D3-D14 ubiquitin ligase in strigolactone signalling. Nature 563, 652–656 (2018).

13. Y. Wang, H. J. Bouwmeester, Structural diversity in the strigolactones. J Exp Bot 69, 2219–2230 (2018).

14. E. B. Aliche, C. Screpanti, A. De Mesmaeker, T. Munnik, H. J. Bouwmeester, Science and application of strigolactones. New Phytol 227, 1001–1011 (2020).

15. L. Wang et al., Transcriptional regulation of strigolactone signalling in Arabidopsis. Nature 583, 277–281 (2020).

16. M. T. Waters, D. C. Nelson, Karrikin perception and signalling. New Phytol 237, 1525–1541 (2023).

17. L. Wang et al., Strigolactone and Karrikin Signaling Pathways Elicit Ubiquitination and Proteolysis of SMXL2 to Regulate Hypocotyl Elongation in Arabidopsis. Plant Cell 32, 2251–2270 (2020).

18. J. P. Stanga, S. M. Smith, W. R. Briggs, D. C. Nelson, SUPPRESSOR OF MORE AXILLARY GROWTH2 controls seed germination and seedling development in Arabidopsis. Plant Physiol 163, 318–330 (2013).

19. I. Soundappan et al., SMAX1-LIKE/D53 Family Members Enable Distinct MAX2-Dependent Responses to Strigolactones and Karrikins in Arabidopsis. Plant Cell 27, 3143–3159 (2015).

20. J. Zheng et al., Karrikin Signaling Acts Parallel to and Additively with Strigolactone Signaling to Regulate Rice Mesocotyl Elongation in Darkness. Plant Cell 32, 2780–2805 (2020).

21. J. C. Beltran, C. Stange, Apocarotenoids: A New Carotenoid-Derived Pathway. Subcell Biochem 79, 239–272 (2016).

22. S. D’Alessandro, Y. Mizokami, B. Legeret, M. Havaux, The Apocarotenoid beta-Cyclocitric Acid Elicits Drought Tolerance in Plants. iScience 19, 461–473 (2019).

23. F. Ramel et al., Carotenoid oxidation products are stress signals that mediate gene responses to singlet oxygen in plants. Proc Natl Acad Sci U S A 109, 5535–5540 (2012).

24. K. P. Jia et al., An alternative, zeaxanthin epoxidase-independent abscisic acid biosynthetic pathway in plants. Mol Plant 15, 151–166 (2022).

25. J. Y. Wang et al., The apocarotenoid metabolite zaxinone regulates growth and strigolactone biosynthesis in rice. Nat Commun 10, 810 (2019).

26. A. Felemban et al., The apocarotenoid beta-ionone regulates the transcriptome of Arabidopsis thaliana and increases its resistance against Botrytis cinerea. Plant J 117, 541–560 (2024).

27. J. Y. Wang et al., Multi-omics approaches explain the growth-promoting effect of the apocarotenoid growth regulator zaxinone in rice. Commun Biol 4, 1222 (2021).

28. A. Ablazov et al., The Apocarotenoid Zaxinone Is a Positive Regulator of Strigolactone and Abscisic Acid Biosynthesis in Arabidopsis Roots. Front Plant Sci 11, 578 (2020).

29. P. Gu et al., The D14-SDEL1-SPX4 cascade integrates the strigolactone and phosphate signalling networks in rice. New Phytol 239, 673–686 (2023).

30. Y. K. Sun et al., Divergent receptor proteins confer responses to different karrikins in two ephemeral weeds. Nature Communications 11, (2020).

31. M. T. Waters et al., A Selaginella moellendorffii Ortholog of KARRIKIN INSENSITIVE2 Functions in Arabidopsis Development but Cannot Mediate Responses to Karrikins or Strigolactones. Plant Cell 27, 1925–1944 (2015).

32. M. Jamil et al., Methyl phenlactonoates are efficient strigolactone analogs with simple structure. Journal of Experimental Botany 69, 2319–2331 (2018).

33. S. A. Stirling et al., Volatile communication in plants relies on a KAI2-mediated signaling pathway. Science 383, 1318–1325 (2024).

34. X. R. Li et al., Nutrient regulation of lipochitooligosaccharide recognition in plants via NSP1 and NSP2. Nat Commun 13, 6421 (2022).

35. K. Mashiguchi et al., Feedback-regulation of strigolactone biosynthetic genes and strigolactone-regulated genes in Arabidopsis. Biosci Biotechnol Biochem 73, 2460–2465 (2009).

36. L. Wang et al., Strigolactone Signaling in Arabidopsis Regulates Shoot Development by Targeting D53-Like SMXL Repressor Proteins for Ubiquitination and Degradation. Plant Cell 27, 3128–3142 (2015).

37. T. Bennett et al., Strigolactone regulates shoot development through a core signalling pathway. Biol Open 5, 1806–1820 (2016).

38. M. Su et al., The LEA protein, ABR, is regulated by ABI5 and involved in dark-induced leaf senescence in Arabidopsis thaliana. Plant Sci 247, 93–103 (2016).

39. M. Mizoguchi et al., Two closely related subclass II SnRK2 protein kinases cooperatively regulate drought-inducible gene expression. Plant Cell Physiol 51, 842–847 (2010).

40. L. C. David et al., N availability modulates the role of NPF3.1, a gibberellin transporter, in GA-mediated phenotypes in Arabidopsis. Planta 244, 1315–1328 (2016).

41. K. H. Ng, H. Yu, T. Ito, AGAMOUS controls GIANT KILLER, a multifunctional chromatin modifier in reproductive organ patterning and differentiation. PLoS Biol 7, e1000251 (2009).

42. M. W. Pfaffl, A new mathematical model for relative quantification in real-time RT-PCR. Nucleic Acids Res 29, e45 (2001).

43. D. Parkhomchuk et al., Transcriptome analysis by strand-specific sequencing of complementary DNA. Nucleic Acids Res 37, e123 (2009).

44. A. Mortazavi, B. A. Williams, K. McCue, L. Schaeffer, B. Wold, Mapping and quantifying mammalian transcriptomes by RNA-Seq. Nat Methods 5, 621–628 (2008).

45. M. Pertea et al., StringTie enables improved reconstruction of a transcriptome from RNA-seq reads. Nat Biotechnol 33, 290–295 (2015).

46. Y. Liao, G. K. Smyth, W. Shi, featureCounts: an efficient general purpose program for assigning sequence reads to genomic features. Bioinformatics 30, 923–930 (2014).

47. M. I. Love, W. Huber, S. Anders, Moderated estimation of fold change and dispersion for RNA-seq data with DESeq2. Genome Biol 15, 550 (2014).

48. D. Veyel et al., PROMIS, global analysis of PROtein-Metabolite Interactions using Size separation in Arabidopsis thaliana. J Biol Chem, (2018).

49. U. S. Hameed et al., Structural basis for specific inhibition of the highly sensitive ShHTL7 receptor. Embo Rep 19, (2018).

50. A. Hungler, A. Momin, K. Diederichs, S. T. Arold, ContaMiner and ContaBase: a webserver and database for early identification of unwantedly crystallized protein contaminants. J Appl Crystallogr 49, 2252–2258 (2016).

51. F. Long, A. A. Vagin, P. Young, G.N. Murshudov, BALBES: a molecular-replacement pipeline. Acta Crystallogr D 64, 125–132 (2008).

52. P. Emsley, K. Cowtan, Coot: model-building tools for molecular graphics. Acta Crystallogr D 60, 2126–2132 (2004).

53. P. V. Afonine et al., Towards automated crystallographic structure refinement with phenix.refine. Acta Crystallogr D 68, 352–367 (2012).

54. J. Mi et al., A manipulation of carotenoid metabolism influence biomass partitioning and fitness in tomato. Metab Eng 70, 166–180 (2022).

55. M. Jamil et al., Striga hermonthica Suicidal Germination Activity of Potent Strigolactone Analogs: Evaluation from Laboratory Bioassays to Field Trials. Plants (Basel) 11, (2022).

56. J. Braguy et al., SeedQuant: a deep learning-based tool for assessing stimulant and inhibitor activity on root parasitic seeds. Plant Physiology 186, 1632–1644 (2021).

