## Supplementary material for "DWARF14 and KARRIKIN INSENSITIVE2 mediate signaling of the apocarotenoid zaxinone in Arabidopsis": All supplemental material including methods, Figures and Tables

**The PDF file includes:**

Materials and Methods

Figs. S1 to S11

Tables S1 to S13

References 28, 37-56

**Other Supplementary Materials for this manuscript include the following:**

Data S1 to S32

**Materials and Methods**

**Plant growth and treatments.** Arabidopsis wild type (*Col-0*), *Atd14-1*, *Atkai2-2*, and *max2-1* mutants (*Col-0* background) were previously described (*37*). Freshly propagated seeds were grown hydroponically in a box system for 2 weeks as described in Ablazov et al. (*28*). Then, plants were transferred to 50 mL black Falcon tubes and grown for 3 weeks (*28*). Five-week old plants were treated with zaxinone (20 µM) or with a mock solution for six hours. Plants were grown in a plant growth chamber (Percival) under controlled conditions (photoperiod of 10/14 h day/night, 22°C, 55% humidity, and 100 µmol m^-2^s^-1^ light intensity).

**Seed germination assays.** Freshly propagated Arabidopsis wild type *Col-0*, *Atd14*, *Atkai2*, and *Atmax2* mutant seeds were surface sterilized with 20% bleach for 15 min and cold-treated for two days in Eppendorf tubes at 4°C and in dark conditions. Seeds were germinated on Petri dishes containing Murashige and Skoog basal salt (MS) medium (1/2 MS salts, 1% agar, 1% sucrose, 0.5 g/L MES, and pH 5.75) with mock, zaxinone (50 µM) or ABA (2 µM). Then Petri dishes were placed on a phytotron with a photoperiod of 16/8 h day/night, 22°C, 55% humidity, and 160 µmol m^-2^s^-1^ light intensity, for four days. Radicle appearance was recorded after 2 days of treatment and seedlings were photographed after 4 days of treatment.

**RNA extraction and gene expression analysis.** Total RNA was extracted from Arabidopsis (WT *Col-0*, *Atd14*, *Atkai2*, and *max2*) roots employing the TRI-Reagent and Direct-zol RNA Miniprep Kit (Zymo Research, R2072) according to the manufacturer’s instructions. The concentration and quality of the RNA samples was determined using a NanoDropTM 2000 Spectrophotometer. For cDNA synthesis, 1 µg of total RNA was used to synthesize total cDNA using the iScript^TM^ cDNA synthesis kit, following the manufacturer’s protocol (Bio-Rad, 1708890). Quantitative real-time PCR analysis (qPCR) was performed using the Advanced^TM^ Universal SYBR Green supermix, according to the manufacturer’s protocol (Bio-Rad, 10000076382). The reaction was prepared in 384-well plates in a total volume of 10 µL. PCR was performed using the StepOnePlus^TM^ Real-Time PCR System. Primer list for ABA, SL and SL-related genes and the normalizer *AtCACS* (AT5G46630) can be found in the Supplementary Table 13 (*28, 38-41*). The expression levels of all analyzed genes was normalized to that of the housekeeping gene *AtCACS* using the equation of 2^-ΔΔCT^ (*42*). Four biological replicates and three technical replicates were used. Data visualization and statistical analysis were performed using the GraphPad Prism 9.1 software.

**RNA sequencing.** RNA sequencing was performed by Novogene (Novogene Co., Ltd., Beijing, China). A total of 2000 ng RNA from Arabidopsis (WT *Col-0*, *Atd14*, *Atkai2* and *max2*) samples treated with mock and zaxinone were sent to Novogene. Messenger RNA was purified from total RNA using poly-T oligo-attached magnetic beads. After fragmentation, the first strand cDNA was synthesized using random hexamer primers, followed by the second strand cDNA synthesis using either dUTP for directional library or dTTP for non-directional library (*43*). The directional library was ready after end repair, A-tailing, adapter ligation, size selection, USER enzyme digestion, amplification, and purification. The library was checked with Qubit and real-time PCR for quantification and bioanalyzer for size distribution detection. Quantified libraries were pooled and sequenced on Illumina platforms, according to effective library concentration and data amount and paired-end reads were generated.

**RNAseq data analysis.** Data analysis was performed by Novogene. Briefly, for quality control, raw data (raw reads) of fastq format were firstly processed through in-house perl scripts. Clean data (clean reads) were obtained by removing reads containing adapter, reads containing ploy-N and low quality reads from raw data. At the same time, Q20, Q30 and GC content the clean data were calculated. All the downstream analyses were based on the clean data with high quality. Reference genome and gene model annotation files were downloaded from genome website (the latest reference genome from TAIR website). Index of the reference genome was built using Hisat2 (*44*) v2.0.5 and paired-end clean reads were aligned to the reference genome using Hisat2 v2.0.5. We selected Hisat2 as the mapping tool for that Hisat2 can generate a database of splice junctions based on the gene model annotation file and thus a better mapping result than other non-splice mapping tools. The mapped reads of each sample were assembled by StringTie (v1.3.3b) (*45*) in a reference-based approach. StringTie uses a novel network flow algorithm as well as an optional de novo assembly step to assemble and quantitate full length transcripts representing multiple splice variants for each gene locus. FeatureCounts (*46*) v1.5.0-p3 was used to count the reads numbers mapped to each gene. And then FPKM of each gene was calculated based on the length of the gene and reads count mapped to this gene. FPKM, expected number of Fragments Per Kilobase of transcript sequence per Millions base pairs sequenced, considers the effect of sequencing depth and gene length for the reads count at the same time, and is currently the most commonly used method for estimating gene expression levels. Differential expression analysis of two conditions/groups (fours biological replicates per condition) was performed using the DESeq2 R package (1.20.0) (*47*). DESeq2 provide statistical routines for determining differential expression in digital gene expression data using a model based on the negative binomial distribution. The resulting p-values were adjusted using the Benjamini and Hochberg's approach for controlling the false discovery rate. Genes with an adjusted p-value (padjust <=0.05) and an absolute fold change of 2 were assigned as differentially expressed.

**Recombinant protein expression and purification.** The expression vectors pGEX-6P-1/*At*D14 and pGEX-6P-1/*At*KAI2 has been described previously 2016 (*6*). *AtDLK2* CDS sequence was amplified from Arabidopsis cDNA (primers DLK2_HISBAMH1 forward ATTGGATCCATGGTGGTTAATCAGAAGATATCCCG and DLK2_HISR NOT1 reverse TTAGCGGCCGCTCAAGGAGGCGCCTCATGACG). Then, it was cloned into the pQLinkH (plasmid # 13667 Addgene) expression vector for further expression and purification in *E. coli*. For GST-tag protein purification, proteins were expressed and purified as previously described in Wang et al. (*25*) with small modifications. Briefly, BL-21 (DE3) *E. coli* cells carrying the pGEX-6P-1(GST-Tag)/*At*D14 or pGEX-6P-1(GST-Tag)/*At*KAI2 were incubated at 37°C until reach an O.D._600_ of 0.6. Then, cultures were induced with 0.5 mM isopropyl-β-D-thiogalactopyranoside (IPTG) and grown at 16°C for 18 hours. Cells were harvested by centrifugation for 10 min at 5000 RPM at 4°C and subsequently resuspended in lysis buffer (50 mM Tris-HCl/pH=8, 200 mM NaCl, and 2 mM dithiothreitol/DTT with detergent Triton X100/0.5%). After sonication on ice for 10 minutes, the lysate was centrifuged for one hour at 17000 RPM at 4°C. The supernatant was incubated with glutathione-sepharose beads (GE Healthcare) for two hours at 4°C. The column was washed three times with lysis buffer for 15 min at 4°C. Afterwards, the GST moiety was cleaved from the protein using PreScission^TM^ Protease (GE Healthcare) at 4°C for two hours. The purified protein was concentrated using 10K Amicon filter units (Merck Millipore).

**Concentration dependence and specificity interaction assays using nanoDSF.** Arabidopsis *At*D14, *At*KAI2, and *At*DLK2 proteins were obtained as described above. A concentration of 6 µM was used for the thermal stability assays. All proteins and metabolites were diluted to the desired concentration in 1x elution buffer. Concentration curves were assayed for *At*D14, *At*KAI2, and *At*DLK2 incubated with zaxinone (1, 10, 50, and 100 µM), (±)-GR24 (1, 50, 100, and 250 µM), (-)-GR25 (0, 50, 100, 250 and 500 µM) and KAR_2_ (1, 50, 100, and 250 µM). Specificity assays (*At*D14 and *At*KAI2) were performed by diluting zaxinone, (±)-GR24, (-)-GR25and KAR_2_ in elution buffer to a final concentration of 50 µM. Proteins were diluted to a concentration of 6 µM. Metabolites were incubated with each protein, immediately loaded into the capillaries, and placed in the Prometheus NT.48 (Nanotemper). Capillaries were run according to Veyel et al. (*48*). All experiments were performed at least twice (*n*=4-5). In all nanoDSF graphs one representative replicate from the unfolding curve for each protein in the absence and presence of the studied metabolite was shown in each graph. Raw data corresponding to the first derivative of the ratio of the fluorescence 350nm/330nm was exported to excel and graphs and statistical analysis were performed using the GraphPad Prism 9.1 software. Experiments were also performed for rice *Os*D14 following the same protocol as for *At*D14.

**Yoshimulactone green (YLG) hydrolysis assay.** *At*D14 protein (3 µM) was mixed with reaction buffer (1x PBS) in a 100 µL volume on a 96-well black plate (Greiner). The fluorescence intensity in the reaction was measured in a SpectraMaxi3 (Molecular Devices) fluorimeter according to Hameed et al.(*49*). *At*D14 was mixed with Zaxinone, and racemic (±)-GR24 using serial dilutions to cover substrates concentration ranges. Protein-substrate mixes were incubated for 30 min. Then, YLG (1 µM; Tokyo Chemical Industry Co. Ltd.) was added and the reaction was incubated for two hours. The change in fluorescence observed over the course of two hours incubation of YLG in buffer without *At*D14 was subtracted from the data collected in the presence of *At*D14. Relative fluorescence was plotted and inhibitory curves and their respective IC_50_ values were calculated using GraphPad Prism 9.1 four-parameter logistic curve. Experiments were also performed for rice *Os*D14 following the same protocol as for *At*D14 but without pre-incubation for (±)-GR24.

**Tryptophan intrinsic fluorescence.** The fluorescence emitted from tryptophans in the structure of *At*D14 (~10 µM) protein was quantified in a flat-bottomed, black 96-well plate (Greiner) using a spectraMaxi3 plate reader (Molecular Devices). In the assay, eight different concentrations (0.0048, 0.8, 1.6, 3.12, 6.25, 12.5, 25, and 50 μM) were used for zaxinone and (±)-GR24. Each reaction was done in triplicate and in a final volume of 50 µL using PBS buffer (100 mM phosphate, pH 6.8, 150 mM NaCl). *At*D14 tryptophans were excited at 280 nm and emission intensity was measured at 333 nm, and the differences in fluorescence intensity were recorded and analyzed. Data were normalized and dissociation coefficient values (*K_d_*) were calculated by fitting to a binding saturation single-site model with GraphPad Prism 9.1 software. Experiments were also performed for rice *Os*D14 following the same protocol as for *At*D14.

**Time-course substrate stability nanoDSF experiments.** *At*D14 protein 6 µM was incubated with reaction buffer (1x PBS), and at each time point (0, 30, 60, 120, 240 min for (±)-GR24; 0, 15, 45, 120, 240 for MP3; 0, 60, 120, 240, 360 min for zaxinone) GR24, MP3, and zaxinone diluted in PBS were added to the protein solution to initiate the reaction. The final concentration of the metabolites was 50 µM for (±)-GR24 and MP3 and 20 µM for zaxinone in the interaction assays with *At*D14. For *At*KAI2 interaction assays (-)-GR24 and zaxinone final concentration was 100 µM. After 4 h ((-)-GR24 and MP3) or 6 h (zaxinone) incubation at 23 °C, all the reactions were loaded in the capillaries (~12 µL) and placed in PrometheusNT.48 (Nanotemper) device. Samples were run and analyzed as described above. Experiments were also performed for rice *Os*D14 following the same protocol as for *At*D14.

***At*D14 protein purification, crystallization and structure determination.** Cells were harvested by centrifugation at 8,000 *g* for 15 mins. Cells from one-liter culture were re-suspended in 25 ml lysis buffer (50 mM Tris-HCl pH 7.5, 250 mM NaCl, 3 mM DTT, 0.2 % Triton X-100) and lysed by sonication. Cell debris was removed by centrifugation at 75,000 *g* for 30 minutes, and proteins were purified from the supernatant using glutathione sepharose 4B resins (GE Healthcare). The N-terminal GST tag of *At*D14 was removed by overnight incubation with Prescission Protease (GE healthcare) at 4°C. After GST cleavage, the resin flow-through containing *At*D14 was further purified on a HiLoad16/60 Superdex 200 prep-grade gel filtration column (GE Healthcare) using a buffer containing 20 mM HEPES pH 7.5, 150 mM NaCl, 3 mM DTT. The purified protein was concentrated at 15 mg/ml and stored at -80°C. 15 mg/ml *At*D14 were first mixed with zaxinone in a 1:5 ratio and then co-crystallized by equilibrating 1.0 μl of protein mixed with 1.0 μl of reservoir solution (0.2 M magnesium chloride hexahydrate, 0.1 M Bis-Tris pH 6.5 and 25 % PEG 3350) using the hanging drop vapor diffusion method. The crystals grew in 7 days at 23°C. For data collection, 25% glycerol was added to the mother liquor as a cryo-protectant, and the crystals were flash-cooled in liquid nitrogen. The data were collected at 100K in the beamline Proxima 1 at the SOLEIL Synchrotron (France), using EIGER-X 16M detector (proposal numbers 20201179 and 20210195). The data were processed in XDS, and checked with the online ContaMiner server for contaminant proteins (*50*). The crystal structure of *At*D14 was determined by molecular replacement using Balbes (*51*) (CCP4 online) with the *At*D14 structure (PDB 4IH4) as a search model. The structure was manually inspected using Coot (*52*) and refined using Phenix Refine (*53*) (Supplementary Table 8). The figures were drawn using PYMOL (pymol.org).

**Modeling of *At*KAI2 structure bound to zaxinone.** The *At*KAI2 model, obtained from the pdb accession 4HTA, was superimposed on the structure of *At*D14 bound to zaxinone using PYMOL. After superimposing the model, the side chains of *At*KAI2 involved in interaction with zaxinone were checked for clashes with the ligand molecule. The figures were drawn using PYMOL.

**Interaction assays with single point amino acid *Atd14* mutants.** We generated *At*D14 (*At*D14^S97A^ and *At*D14^S97C^) point mutants in pGEX-6P-1 by using previously published MBP-tagged *At*D14^S97A^ and *At*D14^S97C^ vectors (*11*). Both mutated versions were expressed and purified as described above. Pure proteins (6 µM) were tested for binding with MP3 and zaxinone at different concentrations (0, 1, 10, 50, 100, and 250 µM). Samples were run and analyzed as described above for the NanoDSF experiments.

**GST Pull-down of C-terminal helix (CTH) of *At*MAX2.** Purified GST-*At*D14 was bound to glutathione sepharose 4B beads. GST-bound *At*D14 was incubated with 100 μM zaxinone for 4 hours first and then 100 μM (±)-GR24 was added to the same sample for competition experiments. In parallel *At*D14 was incubated with 100 μM (±)-GR24 only, and *At*D14 without the addition of GR24 was used as a control. The N-terminally fluorescein isothiocyanate (FITC) labeled CTH sequence of *At*MAX2 (667 – 693) was synthesized by GenScript Biotech Corp. FITC has an excitation wavelength of 488 nm and emits at 520 nm. FITC–CTH was incubated with the above samples for 2 h and then washed 3 times with buffer (50 mM Tris-HCl pH 7.5, 250 mM NaCl, 3 mM DTT). Raisin–associated proteins were eluted with the addition of 20 mM reduced glutathione to the samples. The eluted protein samples were subjected to a Biorad imager using an excitation wavelength of 488 nm.

**Fluorescence anisotropy.** Fluorescence anisotropy assay was carried out on a PHERAstar plate reader (BMG Labtech) with a filter (480/520 nm) and automated polarizer at 25 °C. In the assay, commercially synthesized CTH peptide of AtMAX2 (Genscript Inc.) was labeled with FITC at the N-terminus. 100 μM (±)-GR24 was added to the *At*D14, and the *At*D14 pre-incubated with 100 μM zaxinone for 4 hours respectively, these proteins were then titrated against 50 nM FITC–CTH of *At*MAX2, starting with 250 μM of *At*D14 protein, and then serially diluted until a protein concentration of 490 nM in the buffer containing 50 mM Tris-HCl pH 7.5, 250 mM NaCl, 3 mM DTT. The *K*_d_ value was fitted with one-site specific binding model (Graph Pad Prism).

**ABA quantification.** We perform ultrasound-assisted extraction (UAE) of plant hormones from Arabidopsis roots following the protocol used in Mi et al (*54*). Briefly, for the quantification of endogenous hormone levels 20mg of freeze dried ground root tissues were spike with 2 ng of D_6_-ABA along with 500 µl of 100% MeOH. Sonicate for 15 min follow by centrifuge at 4 °C, 14000 rpm for 8 min. The supernatant is transfer to new 2 ml Eppendorf tube and kept on ice. The extraction was repeated with 500 µl of 100% MeOH without internal standard. Then, the two supernatants were combined into 2 ml Eppendorf tube. Vacuum dry the supernatant collected and stored in -20°C. Hormones were purified using SPE by dissolving the dried sample in 50 µl MeOH, follow by 1ml H_2_O and vortex. C18 SPE column (50mg/ml) was preconditioned with 1 ml MeOH and 1ml H_2_O. Sample solution was loaded onto C18SPE column held on a vacuum manifold. C18 SPE column was washed with 1 ml of H_2_O. Plant hormones were eluted with 1 ml of 50% acetonitrile. Samples were dried in a nitrogen concentrator. Plant hormones were quantified using UHPLC-MS (a Vanquish™ Duo UHPLC Systems coupled with a TSQ Altis™ triple quadrupole mass spectrometer (Thermo Scientific) with a heated-electrospray ionization source) by dissolving plant hormone extracts with 100 µl of 50% acetonitrile and vortexed for 10 s. Sample solution were filtered using a 0.2 µm filter into an amber autosampler vial with an insert. Vials were kept at 4 °C. The chromatographic separation was carried out on an ACQUITY UPLC HSS T3 column (2.1 × 100 mm, 1.8 μm, Waters) and a VanGuard pre-column (2.1 × 5 mm, 1.8 μm, Waters) maintained at 40 °C. Mobile phases consist of 5% aqueous acetonitrile (A) and acetonitrile (B), both containing 0.01% formic acid. Both were employed for eluting ABA with the gradient program: 0-12 min, 5% B to 80% B; 12-13 min, 80% B to 100% B; 13-16 min, 100% B at 0.4 mL/min of flow rate. The MS parameters were as follows: negative ion, 3000 V; sheath gas, 45 Arb; aux gas, 10 Arb; sweep gas, 1 Arb; ion transfer tube temperature, 325 °C; vaporizer temperature, 300 °C; cycle time, 1 s; Q1 resolution (FWHM), 0.7; Q3 resolution (FWHM), 0.7; CID gas (mTorr), 1.5; and chromatographic peak width (sec), 6.

**SL quantification.** Lyophilized Arabidopsis roots were ground to a fine powder. Approximately 20 mg dry weight powder was extracted twice with 1 ml of acetone containing 20 ng GR24 (internal standard) and were sonicated for 15 min at room temperature. After centrifugation, the two extracts were combined and dried under nitrogen. The extract was dissolved in 120 μl of acetonitrile:water (50:50, v:v), followed by filtration using 0.22 μm filter. Detection of MeCLA was performed on UHPLC-Q-Orbitrap-MS (Q-Exactive Plus). Mass Spectrometry parameters were set as follows: capillary temperature of 250 °C, AUX gas temperature of 310 °C, sheath gas of 30 Arb, AUX gas of 10 Arb, spray voltage of 3 kV in positive ion mode, and collision energy of 20 eV in Parallel Reaction Monitoring (PRM) analysis. UHPLC separation was performed on a UHPLC (Thermo ScientificTM UltiMateTM 3000 UHPLC) equipped with an ODS column (Waters C18, 2.1 × 100 mm, 1.7 μm). The column temperature was maintained at 30 °C. The mobile phase consisted of water:methanol 25:75(v/v) (solvent A) and methanol (solvent B), both of which contained 0.1% [v/v] formic acid. LC separation was conducted with a multi-step gradient at a flow rate of 0.2 ml/min. 20% B to 45% B in 5min; 55% B at 10 min; 85% B at 15 min; 100% B at 17 min, hold at 100% B for 4 min and then equilibrate at 20% for 4 min. Quantification of MeCLA in Arabidopsis was performed with PRM of ion pairs for GR24 and MeCLA using the following mass transitions: GR24 299.0914 > 97.0288; MeCLA 347.18530 > 97.02870.

**Striga bioassays.** Arabidopsis WT and mutant plants treated with mock and zaxinone were compared for their strigolactone producing capacity through parasitic seeds germination bioassays by adopting a procedure described by Jamil et al. (*55*). For the purpose, the Arabidopsis plants were grown hydroponically in boxes and 50 ml tubes for two and three weeks, respectively, under +Pi (four weeks) and –Pi (1 week) conditions. The strigolactones were collected from the root exudates of each treated plant through C18 column which were applied on pre-conditioned *Striga* seeds to see effect on germination. For *col-0* and *Atd14* bioassays experiment we combined together two 50 mL Falcon tubes while for the *Col-0*, *kai2* and *max2* experiment, we used one 50 mL Falcon tube to collect the exudates. For pre-conditioning, the *Striga hermonthica* seeds were surface sterilized with 50% commercial bleach for five minutes and washed with sterilized milliQ water six times. The seeds were dried in a laminar flow cabinet and spread uniformly on 9 mm glass fiber filter paper discs (~50-100 seeds per disc). Then, 12 discs with *Striga* seeds were transferred into a Petri plate, containing a Whatman filter paper and moistened with 3 ml sterilized water. The Petri plates were sealed and wrapped in aluminum foil and incubated at 30 °C for 10 days. The pre-conditioned seeds were treated with each sample (at 55 µl per disc; *n*=4 for each treatment) and incubated again at 30 °C for 24 h. The seeds were scanned by a microscope and germinated and total *Striga* seeds were counted using the software SeedQuant (*56*) to calculate the germination percentage.

**Statistical analyses.** All experiments were performed with at least 3 biological replicates. Statistical tests were carried out either by ANOVA, non-paired two-tails Student *t*-test, binding saturation single-site model and four-parameter logistic curve using the software GraphPad Prism 9.1.0. For differential gene expression analysis the package DESeq2 on R was used.

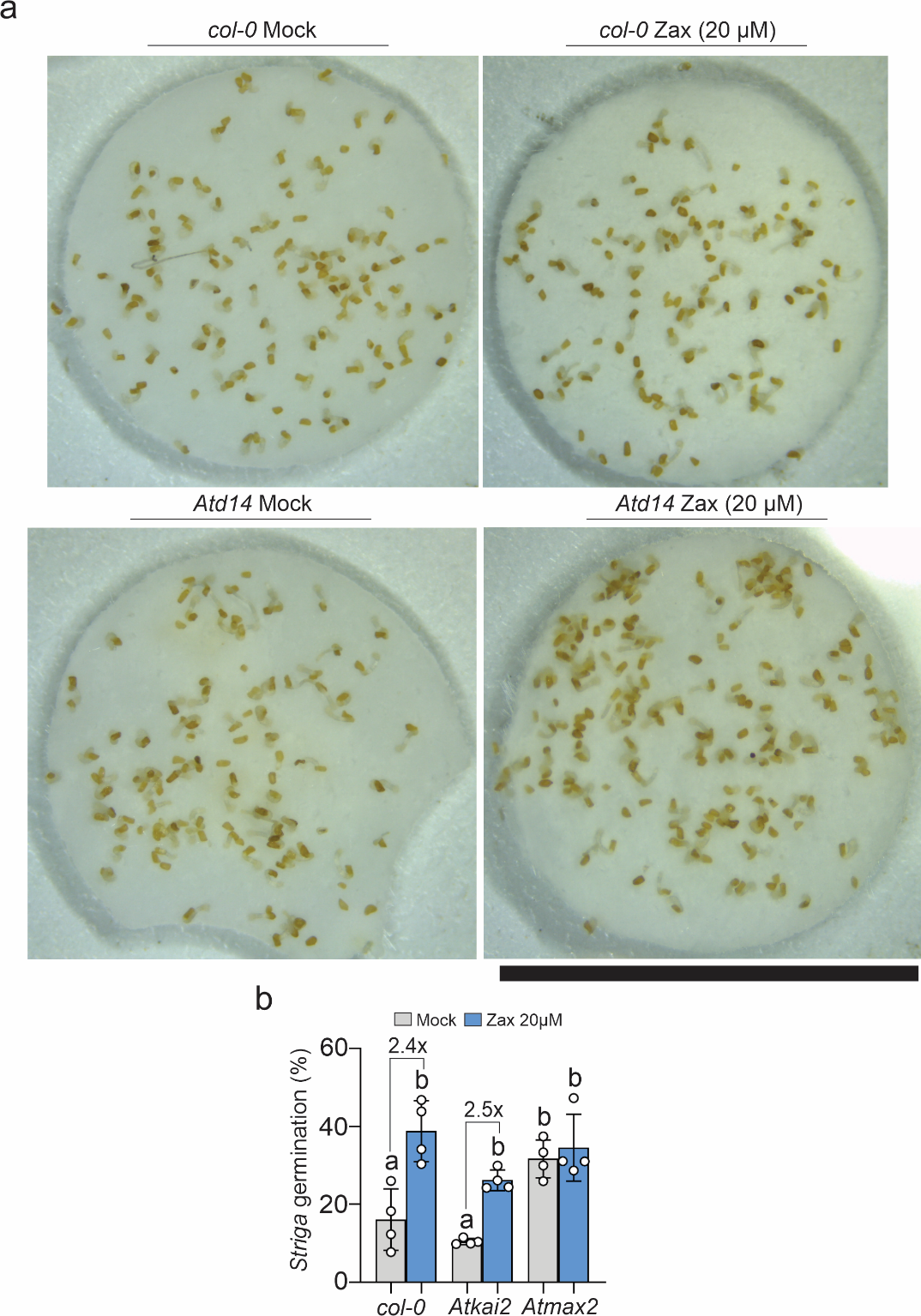

**Figure S1. Seed germination assays. a** Striga germination bioassay using the root exudates from 5-weeks old wt *col-0* and mutant *Atd14* Arabidopsis plants treated with mock (grey) or zaxinone (Zax) (blue; 20 µM) for 6 hours. The photos of the disc are representative images from the experiment. **b** Striga germination bioassay using the root exudates from 5-weeks old *col-0*, *Atkai2*, and *Atmax2* mutant seeds of Arabidopsis plants treated with mock (grey) or zaxinone (Zax) (blue; 20 µM) for 6 hours. One-way ANOVA with multiple comparisons was performed to assign statistical significance to each genotype and treatment (*n*=4). Letters denote significance differences (p<0.05). Photos in (a) were used to do the zoom in to build Figure 1c. Scale bar: 3 mm.

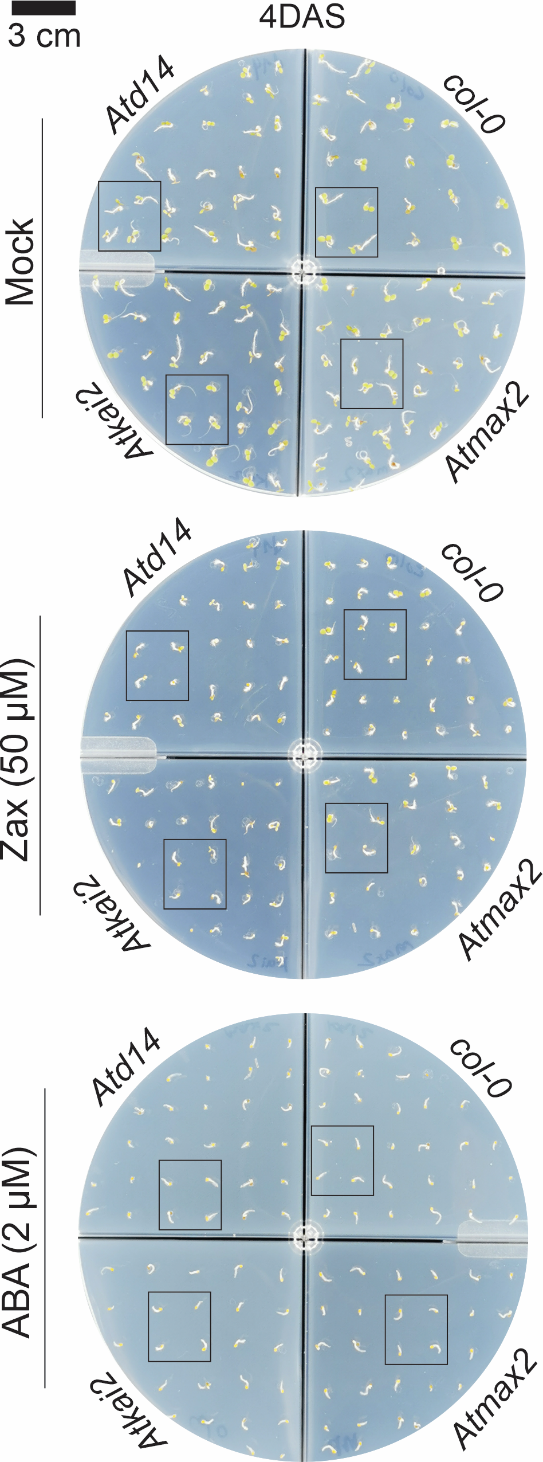

**Figure S2. Seed germination assays.** Germination assay for *col-0*, *Atd14*, *Atkai2*, and *Atmax2* mutant seeds on plates supplemented with mock (1% DMSO), zaxinone (Zax) (50 µM), and ABA (2 µM). Phenotype was recorded at four days after sowing (DAS). The black squares indicate the region taken for the zoom in shown in the main figure 1d.

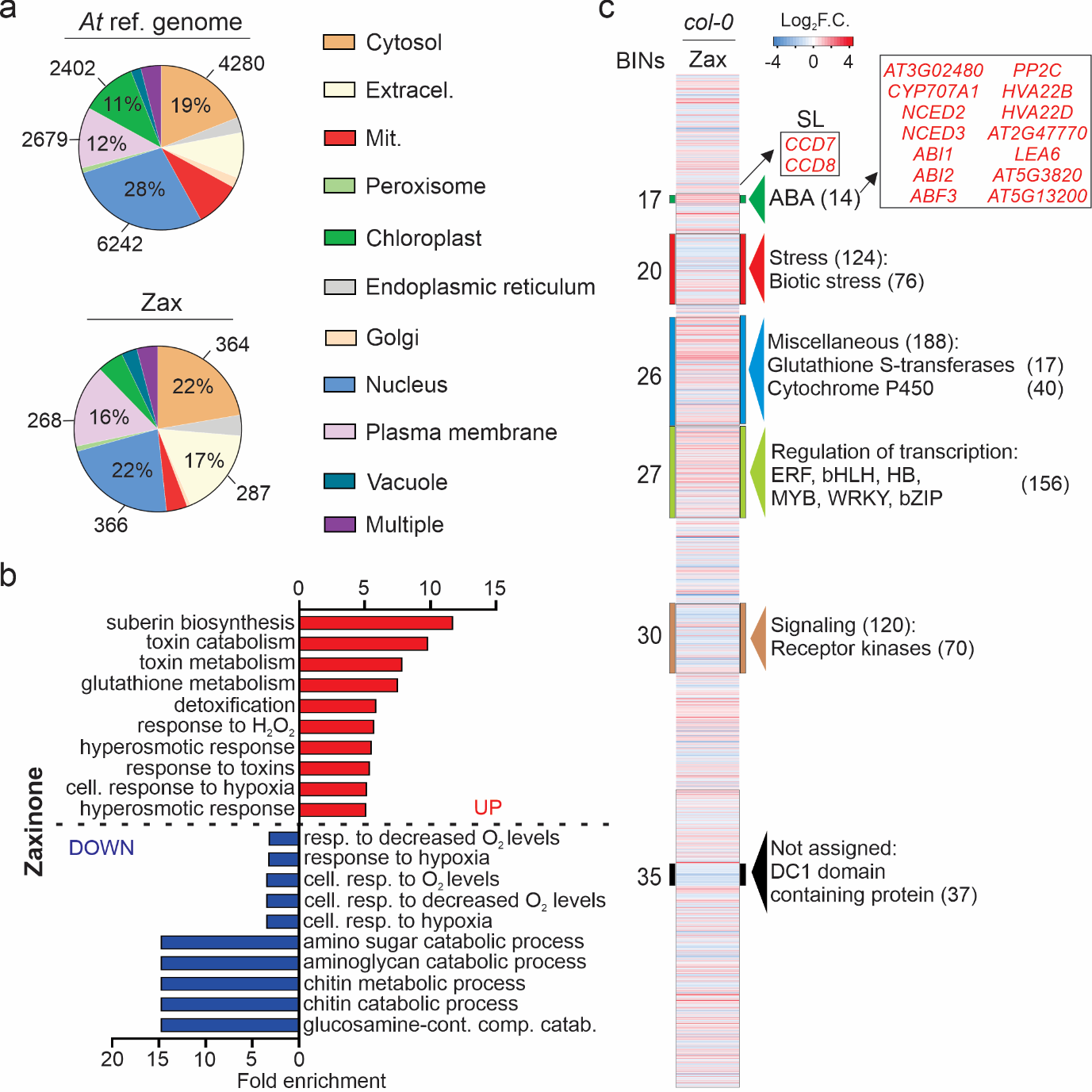

**Figure S3. Subcellular localization and GO analysis of DEGs identified upon zaxinone treatment. a** Distribution of cellular compartments for genes present in the reference Arabidopsis genome (TAIR) and differentially expressed genes (DEGs) in the zaxinone treatment. Subcellular localization analysis was performed in SUBA4 online software (https://suba.live/). Genes encoding proteins with 2 or more localizations were grouped in the “multiple” category. All selected DEGs fulfill the threshold Log2 F.C.>1 (up) or Log2 F.C.< -1 (down) and padj<0.05. **b** Gene ontology (GO) enrichment analysis of up- (red) and downregulated (blue) genes identified in the RNAseq analysis of Arabidopsis roots exposed to Mock or zaxinone treatments. Fold enrichment analysis of GO terms related to biological process showing the top 10 enriched processes in Arabidopsis roots treated with zaxinone (20 µM). A fold enrichment analysis was performed using a Fisher’s exact test with FDR correction (*p*<0.05) using the PANTHER overrepresentation analysis (Gene Ontology database released on 18.08.2021). **c** Heatmap representation showing significantly enriched molecular functions (p<0.05) annotated with MapMan (heatmap scale is Log_2_ fold). The RNAseq experiment was performed by applying 20 µM zaxinone to the hydroponic media of 5-weeks old Arabidopsis plants for 6 hours.

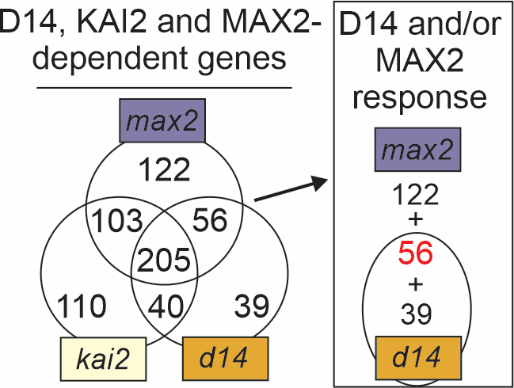

**Figure S4. D14. D14-, KAI2- and MAX2-dependent genes.** Venn diagram showing the number of genes that require D14, KAI2 and/or MAX2 to modulate their expression. The RNAseq data obtained in the *d14*, *kai2* and *max2* mutants treated with mock and zaxinone was used to build the Venn diagram.

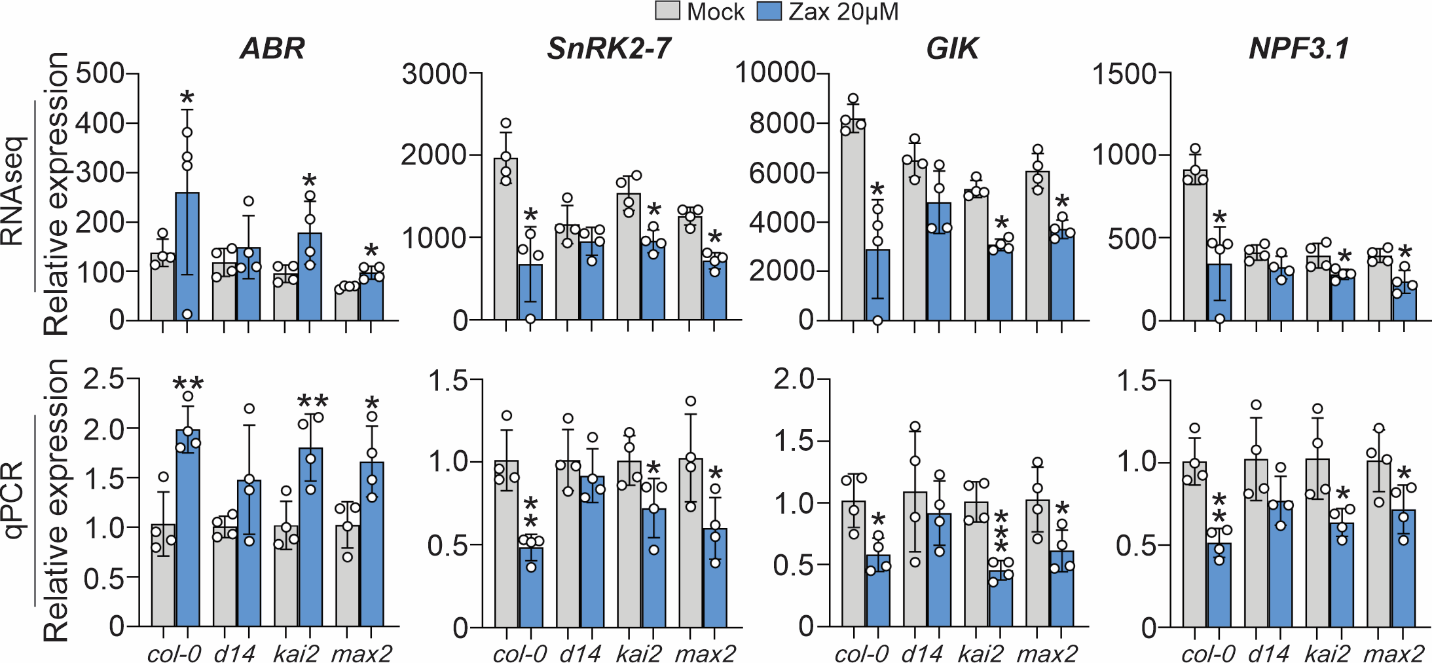

**Figure S5. Zaxinone-triggered transcriptional response *via At*D14.** RNAseq expression profile of *ABR*, *SnRK2-7*, *GIK* and *NPF3.1* genes and their validation by qPCR. These genes correspond to a subset of genes that only require *At*D14 (MAX2-independent response) to control their expression. For qPCR a non-paired two tails *t*-test was performed to determine significance. *: 0.05, **: 0.005, ***: 0.001. DEGs fulfill the threshold Log_2_ F.C.>1 (up) or Log_2_ F.C.< -1 (down) and padj<0.05. The RNAseq experiment was performed by applying 20 µM zaxinone (Zax) to the hydroponic media of 5-weeks old Arabidopsis plants for 6 hours.

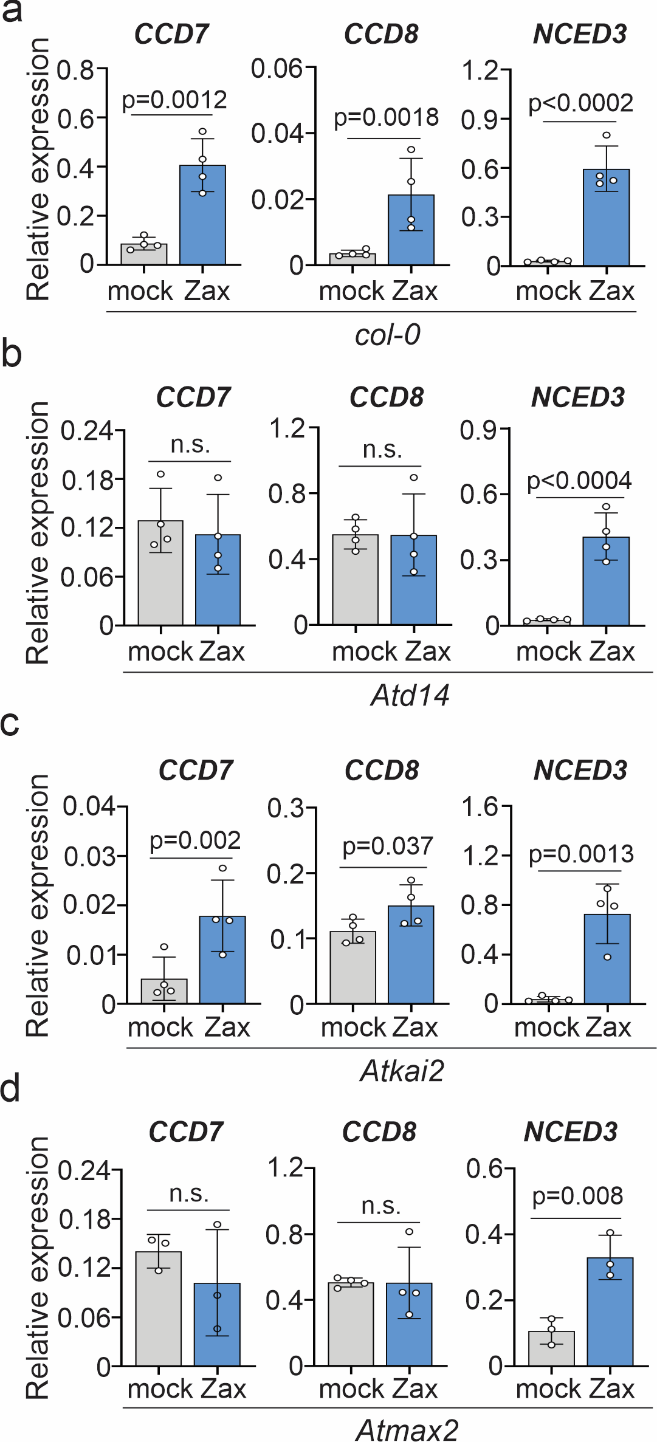

**Figure S6. Gene expression quantification of SL and ABA biosynthetic genes by qPCR. a**-**d** Transcript abundance of SLs and ABA-responsive genes *CCD7*, *CCD8*, and *NCED3* in col-0 (**a**), *Atd14* (**b**), *Atkai2* (**c**), and *Atmax2* (**d**) backgrounds treated with zaxinone (20 µM) or mock, for six hours and measured by qPCR. *CACS* gene was used as normalizer. The data are the mean ± SE of 4 biological replicates and 3 technical replicates. Unpaired two tails Student’s *t*-test was performed to assess significance. The experiment was performed by applying 20 µM zaxinone (Zax) to the hydroponic media of 5-weeks old Arabidopsis *col-0* and mutants for 6 hours.

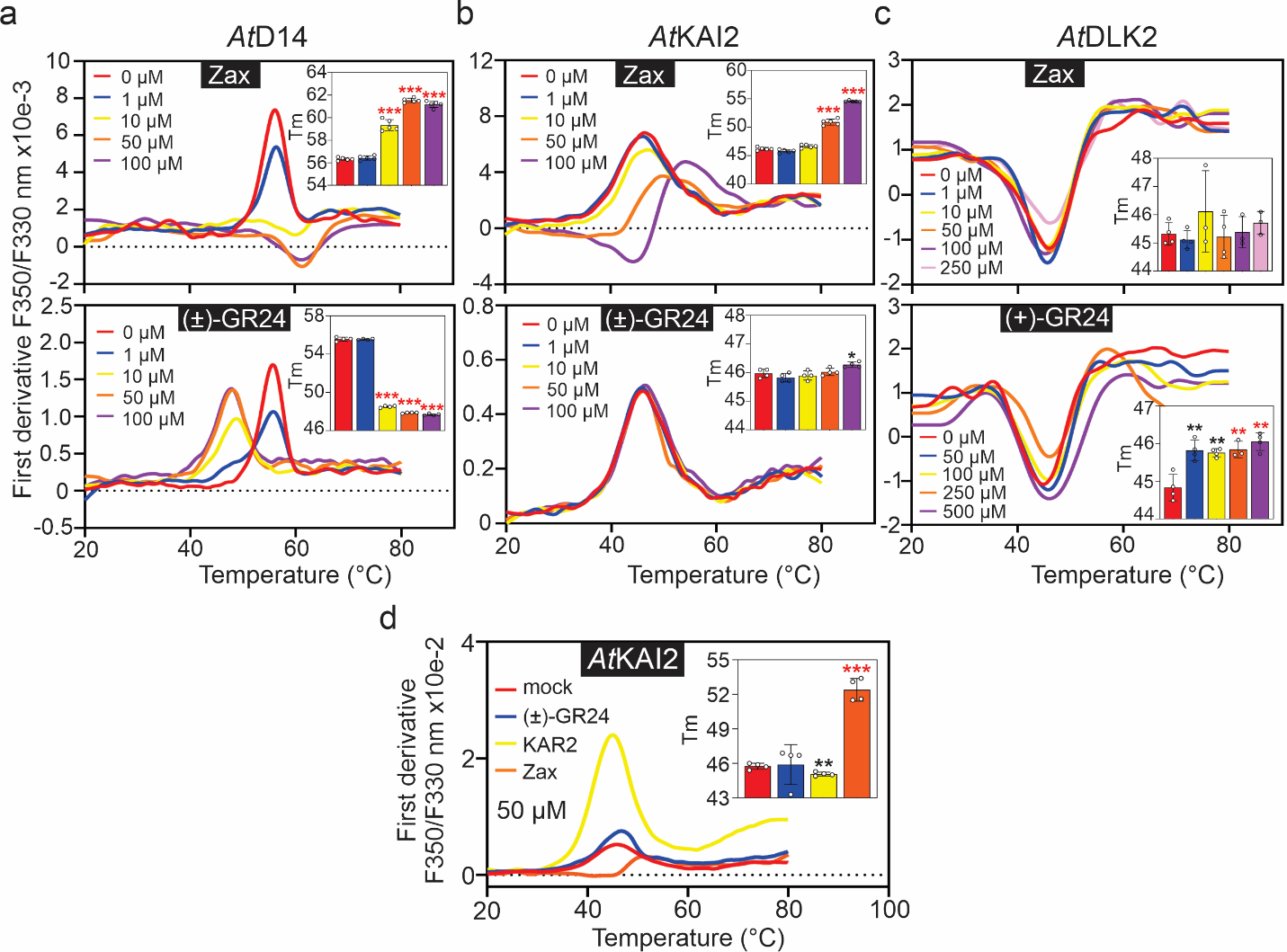

**Figure S7. Thermal shift assays. a-d** Nano differential scanning fluorimetry (nanoDSF) assays. **a-b** NanoDSF curves of *At*D14 and *At*KAI2 in the absence (mock) and presence of zaxinone (0-100 µM) and racemic (±)-GR24 (0-100 µM). **c** NanoDSF curves of *At*DLK2 in the absence and presence of increasing zaxinone and (+)-GR24. **d** NanoDSF assay for *At*KAI2 in the absence and presence of (±)-GR24, KAR2, and zaxinone. T_m_ values were calculated using default settings in PrometheusNT.48 software (means ± SE, *n=4*). * p<0.05; ** p<0.005; *** p<0.0005 (unpaired two tails Student *t*-test). P-value (<0.05) and a shift in the melting profile of at least 1°C were used to define binding (red asterisks). A protein concentration of ~7 µM was used for all proteins.

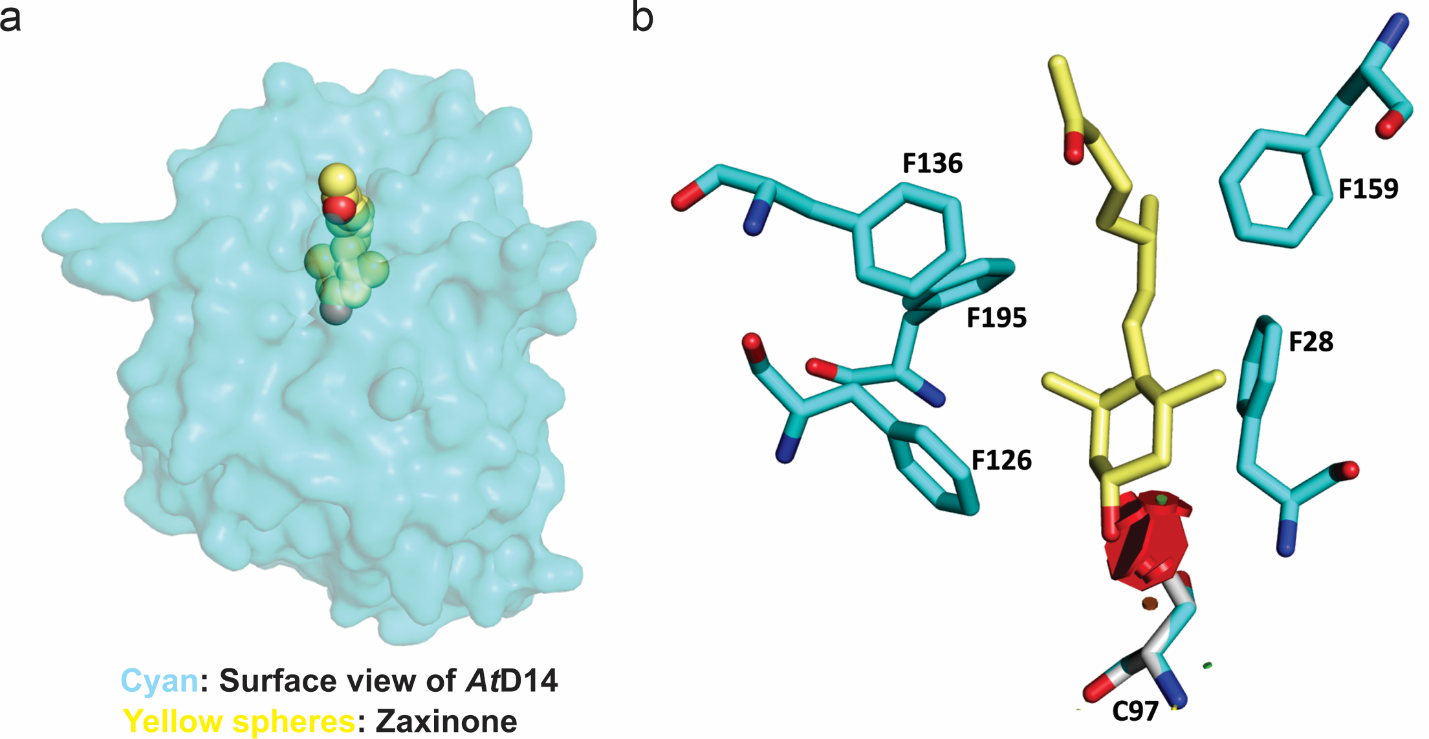

**Figure S8. Structural analysis of *At*D14-zaxinone binding. a** Surface view of the zaxinone (yellow and red spheres) bound to *AtD14* (cyan) at its binding pocket. **b** Zaxinone (yellow sticks) and amino acids of *At*D14 involved in the binding (cyan sticks) where C97 (gray stick) mutation resulted in strong clashes (big red dots) between the sulfhydryl group of cysteine and hydroxyl group of zaxinone.

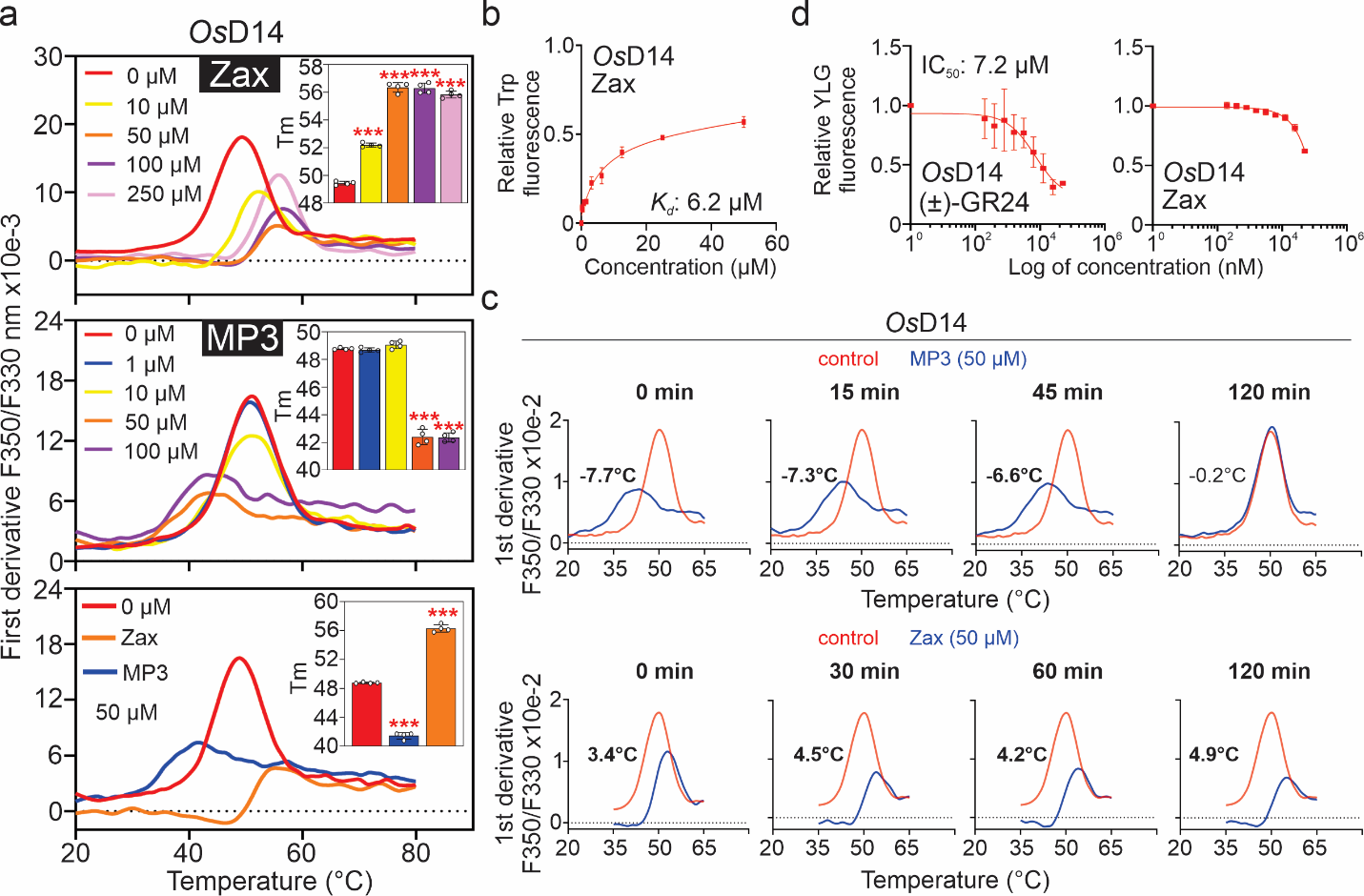

**Figure S9. *In vitro* assays with recombinant *Os*D14 and zaxinone. a** NanoDSF curves of *Os*D14 (6 µM) in the absence and presence of zaxinone (0-250 µM) and MP3 (0-100 µM). Melting curve and *T_m_* shift calculation for the *Os*D14 interaction with MP3 (50 µM) and zaxinone (50 µM). **b** Tryptophan fluorescence assay for *Os*D14 (10 µM) and zaxinone interaction. **c** Melting temperature curves of *Os*D14 with or without pre-incubation with MP3 (50 µM) and zaxinone (50 µM) for the indicated time period. Differences in melting temperature between the pre-incubated and non-treated protein are shown for each graph. *T_m_* values were calculated using default settings in PrometheusNT.48 software (means ± SE, *n=4*; unpaired two tails Student *t*-test). P-value (<0.05) and a shift in the melting profile of at least 1°C were used to define binding (bold-faced temperature values). **d** YLG hydrolysis by *Os*D14 (6 µM) in the presence of increasing (±)*-*GR24 and zaxinone concentrations. Data are the mean ± SE, *n=3*.

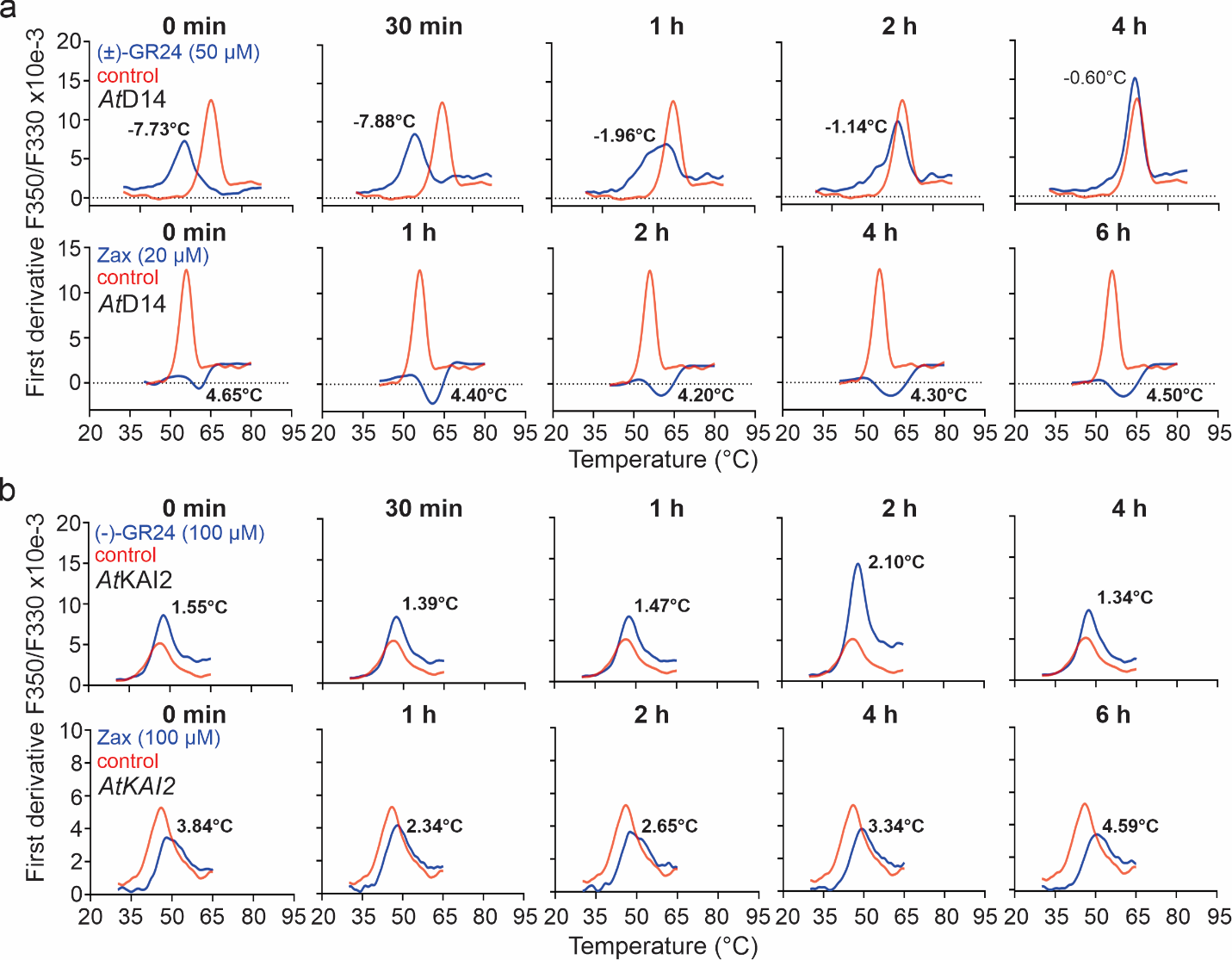

**Figure S10. *At*D14 and *At*KAI2 time point nanoDSF experiments with (±)-GR24 and zaxinone. a-b** Melting temperature curves of *At*D14 (a) and *At*KAI2 (b) with or without pre-incubation with (±)*-*GR24 or (-)-GR24 and zaxinone for the indicated time period. Differences in melting temperature between the pre-incubated and non-treated protein are shown for each graph. Tm values were calculated using default settings in PrometheusNT.48 software (means ± SE, *n=4*; unpaired two tails Student *t*-test). P-value (<0.05) and a shift in the melting profile of at least 1°C were used to define binding (bold-faced temperature values). A protein concentration of ~7 µM was used for all proteins.

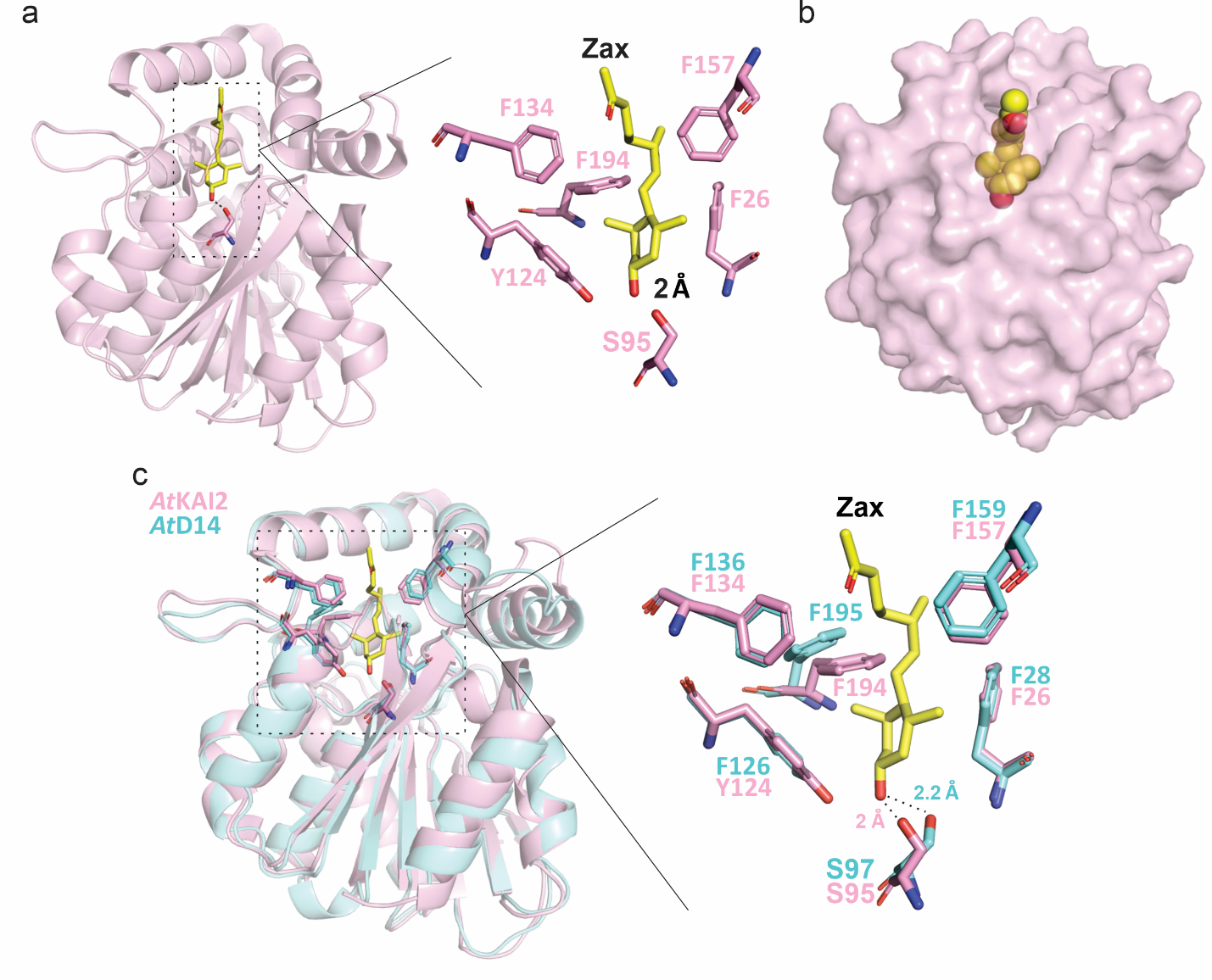

**Figure S11. *In silico* modelling representing the binding between zaxinone and *At*KAI2. a** *In silico* modelling of zaxinone bound to *At*KAI2. Zaxinone (yellow sticks) interacts with *At*KAI2 protein (pink) via six amino acid residues in its binding site (F134, S95, F157, F194, F26 and Y124). The predicted interaction has a resolution of 2 Å and occurs specifically between the OH group of zaxinone and the OH group of the serine 95 (S95) in the *At*KAI2 protein (right side zoom image). **b** Surface view of *At*KAI2 and zaxinone tightly bound in its binding pocket. **c** *In-silico* overlapping between *At*KAI2 (pink) and *At*D14 (cyan) bound to zaxinone.

**Table S1**. NanoDSF specificity experiments for *At*D14 and *A*tKAI2 (data used for the preparation of Figure 3a and Supplementary Fig. 7d).

|  |  | *At*D14 | | | *At*KAI2 | | |
| --- | --- | --- | --- | --- | --- | --- | --- |
| **Apocarotenoid** | **Concentration** **(μM)** | **T_m_ (°C)** | **ΔT_m_** | **p-value** | **T_m_ (°C)** | **ΔT_m_** | **p-value** |
| None | 0 | 55.50 |  |  | 45.73 |  |  |
| (±)-GR24 | 50 | 47.36 | **-8.14** | **<0.0001** | 45.89 | 0.165 | 0.87 |
| Zaxinone | 50 | 60.23 | **4.73** | **<0.0001** | 52.39 | **6.663** | **<0.0001** |
| KAR_2_ | 50 | 55.02 | -0.48 | **0.0001** | 45.07 | -0.66 | **0.0052** |

*Negative T_m_ values represent destabilization of the protein, while positive values represent stabilization. A two tail non-paired student *t*-test was performed to compare control (protein plus elution buffer) and treatment with zaxinone (*n*=4). P-value (≤ 0.05) and a shift in temperature of at least 1 °C were used to define binding (red bold-faced values). This applies for all nanoDSF experiments shown below.

**Table S2**. NanoDSF curve associated to the interaction between zaxinone and *At*D14 (data for preparation of Supplementary Figure 7a upper panel).

|  |  | *At*D14 | | |
| --- | --- | --- | --- | --- |
| **Apocarotenoid** | **Concentration** **(μM)** | **T_m_ (°C)** | **ΔT_m_** | **p-value** |
| Zaxinone | 0 | 56.34 |  |  |
|  | 1 | 56.43 | 0.09 | 0.44 |
|  | 10 | 59.33 | **2.99** | **<0.0001** |
|  | 50 | 61.54 | **5.20** | **<0.0001** |
|  | 100 | 61.18 | **4.84** | **<0.0001** |

**Table S3**. NanoDSF curve associated to the interaction between GR24 and *At*D14 (data for preparation of Supplementary Figure 7a bottom panel).

|  |  | *At*D14 | | |
| --- | --- | --- | --- | --- |
| **Metabolite** | **Concentration** **(μM)** | **T_m_ (°C)** | **ΔT_m_** | **p-value** |
| (±)-GR24 | 0 | 55.57 |  |  |
|  | 1 | 55.54 | -0.035 | 0.82 |
|  | 10 | 48.53 | **-7.040** | **<0.0001** |
|  | 50 | 47.89 | **-7.686** | **<0.0001** |
|  | 100 | 47.66 | **-7.910** | **<0.0001** |

**Table S4**. NanoDSF curve associated to the interaction between zaxinone and *At*KAI2 (data for preparation of Supplementary Figure 7b upper panel).

|  |  | *At*KAI2 | | |
| --- | --- | --- | --- | --- |
| **Apocarotenoid** | **Concentration** **(μM)** | **T_m_ (°C)** | **ΔT_m_** | **p-value** |
| Zaxinone | 0 | 46.22 |  |  |
|  | 1 | 45.81 | -0.41 | **0.05** |
|  | 10 | 46.65 | 0.44 | **0.04** |
|  | 50 | 50.90 | **4.68** | **<0.0001** |
|  | 100 | 54.55 | **8.34** | **<0.0001** |

**Table S5**. NanoDSF curve associated to the interaction between GR24 and *At*KAI2 (data used for the preparation of Supplementary Figure 7b bottom panel).

|  |  | *At*KAI2 | | |
| --- | --- | --- | --- | --- |
| **Metabolite** | **Concentration** **(μM)** | **T_m_ (°C)** | **ΔT_m_** | **p-value** |
| (±)-GR24 | 0 | 45.97 |  |  |
|  | 1 | 45.82 | 0.085 | 0.20 |
|  | 10 | 45.90 | 0.213 | 0.52 |
|  | 50 | 46.03 | 0.06 | 0.62 |
|  | 100 | 46.27 | 0.3 | **0.01** |

**Table S6**. NanoDSF curve associated to the interaction between zaxinone and *At*DLK2 (data used for the preparation of Supplementary Figure 7c upper panel).

|  |  | *At*DLK2 | | |
| --- | --- | --- | --- | --- |
| **Apocarotenoid** | **Concentration** **(μM)** | **T_m_ (°C)** | **ΔT_m_** | **p-value** |
| Zaxinone | 0 | 45.32 |  |  |
|  | 1 | 45.12 | -0.206 | 0.44 |
|  | 10 | 46.12 | 0.795 | 0.33 |
|  | 50 | 45.22 | -0.103 | 0.81 |
|  | 100 | 45.38 | 0.061 | 0.87 |
|  | 250 | 45.57 | 0.247 | 0.26 |

**Table S7**. NanoDSF curve associated to the interaction between GR24*^ent^*^-5DS^ and *At* DLK2 (data used for the preparation of Supplementary Figure 7c bottom panel).

|  |  | *At*DLK2 | | |
| --- | --- | --- | --- | --- |
| **Metabolite** | **Concentration** **(μM)** | **T_m_ (°C)** | **ΔT_m_** | **p-value** |
| (-)-GR2 | 0 | 44.83 |  |  |
|  | 50 | 45.82 | 0.99 | 0.05 |
|  | 100 | 45.76 | 0.93 | <0.0001 |
|  | 250 | 45.85 | **1.02** | **<0.0001** |
|  | 500 | 46.06 | **1.23** | **<0.0001** |

**Table S8.** Data collection and refinement statistics of *At*D14-zaxinone crystal structure.

| **Parameters** | ***At*D14-Zaxinone** |
| --- | --- |
| Resolution range | 44.87 - 2.4 (2.486 - 2.4) |
| Space group | P 1 21 1 |
| Unit cell | 43.82 69.58 89.86 90 92.85 90 |
| Total reflections | 138327 (8268) |
| Unique reflections | 20924 (1788) |
| Multiplicity | 6.6 (4.6) |
| Completeness (%) | 98.49 (85.84) |
| Mean I/sigma(I) | 17.77 (2.77) |
| Wilson B-factor | 44.66 |
| R-merge | 0.07363 (0.5171) |
| R-meas | 0.07989 (0.5824) |
| R-pim | 0.03065 (0.2601) |
| CC1/2 | 0.999 (0.845) |
| CC* | 1 (0.957) |
| Reflections used in refinement | 20920 (1788) |
| Reflections used for R-free | 1046 (90) |
| R-work | 0.2062 (0.2817) |
| R-free | 0.2369 (0.3277) |
| CC(work) | 0.940 (0.716) |
| CC(free) | 0.924 (0.559) |
| Number of non-hydrogen atoms | 4192 |
| macromolecules | 4087 |
| ligands | 40 |
| solvent | 65 |
| Protein residues | 523 |
| RMS(bonds) | 0.008 |
| RMS(angles) | 1.19 |
| Ramachandran favored (%) | 97.29 |
| Ramachandran allowed (%) | 2.32 |
| Ramachandran outliers (%) | 0 |
| Clashscore | 2.94 |
| Average B-factor | 32.74 |
| macromolecules | 32.29 |
| ligands | 59.89 |
| solvent | 43.99 |
| Number of TLS groups | 3 |

Statistics for the highest-resolution shell are shown in parentheses.

**Table S9**. NanoDSF time course experiments for *At*D14 and (±)-GR24 (data used for the preparation of Figure 3h and Supplementary Figure 10a).

|  |  |  | *At*D14 | | |
| --- | --- | --- | --- | --- | --- |
| **Metabolite** | **Concentration** **(μM)** | **Time point (h)** | **T_m_ (°C)** | **ΔT_m_** | **p-value** |
| GR24 | 0 | 0 | 55.57 |  |  |
|  | 50 | 0 | 48.1 | **-7.73** | **<0.0001** |
|  | 50 | ½ | 47.9 | **-7.88** | **<0.0001** |
|  | 50 | 1 | 53.8 | **-1.96** | **<0.0001** |
|  | 50 | 2 | 54.7 | **-1.14** | **<0.0001** |
|  | 50 | 4 | 55.2 | -0.60 | <0.0001 |

**Table S10**. NanoDSF time course experiments for *At*D14 and zaxinone (data used for the preparation of Figure 3h and Supplementary Figure 10a).

|  |  |  | *At*D14 | | |
| --- | --- | --- | --- | --- | --- |
| **Apocarotenoid** | **Concentration** **(μM)** | **Time point (h)** | **T_m_ (°C)** | **ΔT_m_** | **p-value** |
| Zax | 0 | 0 | 55.57 |  |  |
|  | 20 | 0 | 60.40 | **4.65** | **<0.0001** |
|  | 20 | 1 | 60.20 | **4.36** | **<0.0001** |
|  | 20 | 2 | 60.05 | **4.20** | **<0.0001** |
|  | 20 | 4 | 60.12 | **4.27** | **<0.0001** |
|  | 20 | 6 | 60.31 | **4.47** | **<0.0001** |

**Table S11**. NanoDSF time course experiments for *At*D14 and (-)-GR24 (data used for the preparation of Figure 3h and Supplementary Figure 10b).

|  |  |  | *At*KAI2 | | |
| --- | --- | --- | --- | --- | --- |
| **Metabolite** | **Concentration** **(μM)** | **Time point (h)** | **T_m_ (°C)** | **ΔT_m_** | **p-value** |
| (-)-GR24 | 0 | 0 | 46.04 |  |  |
|  | 100 | 0 | 47.58 | **1.55** | **<0.0001** |
|  | 100 | ½ | 47.43 | **1.39** | **<0.0001** |
|  | 100 | 1 | 47.51 | **1.47** | **<0.0001** |
|  | 100 | 2 | 48.14 | **2.10** | **<0.0001** |
|  | 100 | 4 | 47.38 | **1.34** | **<0.0001** |

**Table S12**. NanoDSF time course experiments for *At*D14 and zaxinone (data used for the preparation of Figure 3h and Supplementary Figure 10b).

|  |  |  | *At*KAI2 | | |
| --- | --- | --- | --- | --- | --- |
| **Apocarotenoid** | **Concentration** **(μM)** | **Time point (h)** | **T*m* (°C)** | **ΔT_m_** | **p-value** |
| Zax | 0 | 0 | 45.79 |  |  |
|  | 100 | 0 | 49.64 | 3.84 | **<0.0001** |
|  | 100 | 1 | 48.14 | 2.34 | **<0.0001** |
|  | 100 | 2 | 48.44 | 2.65 | **<0.0001** |
|  | 100 | 4 | 49.13 | 3.34 | **<0.0001** |
|  | 100 | 6 | 50.38 | 4.59 | **<0.0001** |

**Table S13.** List of primers for qPCR used in this work.

| **Gene** | **Primer F’** | **Primer R’** | **Reference** |
| --- | --- | --- | --- |
| *AtMAX3* | TGGCGACGACAAACTACTCC | TGTTCCACCCGTTTAGAGGC | Ablazov et al., 2020 |
| *AtMAX4* | GTTTTACCCGATGCTAGGATC | TGATGCTGCACATATCCATCG |  |
| *AtNCED3* | AGAGCCTTTACATCTCAAAAT | AGACGATAATGGCGGCTGAG |  |
| *AtABR* | AGACTCCTTGCAACAGACTGGAC | TCCTTGACGACATCAGCAGCTC | Su et al., 2016 |
| *AtSnRK2.7* | GAGAATAACTGTACCGGAAATCGAA | CGGCGGCACAACCAAA | Mizoguchi et al., 2010 |
| *AtGIK* | GTATCTCCACTTACGCTCGTCGG | GTACCCCGCAGAGTCACAACAG | Ng et al., 2009 |
| *AtNPF3.1* | TCGGCGGTTTTCAACTTTAC | TTACTATTGCCGCCTTGTCC | David et al., 2016 |
| *AtCACS* | ACTCAGGAAGGTGTACGGTCA | TGCATTTGGAACAGGTTTGT | Ablazov et al., 2020 |

**References**

28.	A. Ablazov *et al.*, The Apocarotenoid Zaxinone Is a Positive Regulator of Strigolactone and Abscisic Acid Biosynthesis in Arabidopsis Roots. *Front Plant Sci* **11**, 578 (2020).

37.	T. Bennett *et al.*, Strigolactone regulates shoot development through a core signalling pathway. *Biol Open* **5**, 1806-1820 (2016).

38.	M. Su *et al.*, The LEA protein, ABR, is regulated by ABI5 and involved in dark-induced leaf senescence in Arabidopsis thaliana. *Plant Sci* **247**, 93-103 (2016).

39.	M. Mizoguchi *et al.*, Two closely related subclass II SnRK2 protein kinases cooperatively regulate drought-inducible gene expression. *Plant Cell Physiol* **51**, 842-847 (2010).

40.	L. C. David *et al.*, N availability modulates the role of NPF3.1, a gibberellin transporter, in GA-mediated phenotypes in Arabidopsis. *Planta* **244**, 1315-1328 (2016).

41.	K. H. Ng, H. Yu, T. Ito, AGAMOUS controls GIANT KILLER, a multifunctional chromatin modifier in reproductive organ patterning and differentiation. *PLoS Biol* **7**, e1000251 (2009).

42.	M. W. Pfaffl, A new mathematical model for relative quantification in real-time RT-PCR. *Nucleic Acids Res* **29**, e45 (2001).

43.	D. Parkhomchuk *et al.*, Transcriptome analysis by strand-specific sequencing of complementary DNA. *Nucleic Acids Res* **37**, e123 (2009).

44.	A. Mortazavi, B. A. Williams, K. McCue, L. Schaeffer, B. Wold, Mapping and quantifying mammalian transcriptomes by RNA-Seq. *Nat Methods* **5**, 621-628 (2008).

45.	M. Pertea *et al.*, StringTie enables improved reconstruction of a transcriptome from RNA-seq reads. *Nat Biotechnol* **33**, 290-295 (2015).

46.	Y. Liao, G. K. Smyth, W. Shi, featureCounts: an efficient general purpose program for assigning sequence reads to genomic features. *Bioinformatics* **30**, 923-930 (2014).

47.	M. I. Love, W. Huber, S. Anders, Moderated estimation of fold change and dispersion for RNA-seq data with DESeq2. *Genome Biol* **15**, 550 (2014).

48.	D. Veyel *et al.*, PROMIS, global analysis of PROtein-Metabolite Interactions using Size separation in Arabidopsis thaliana. *J Biol Chem*, (2018).

49.	U. S. Hameed *et al.*, Structural basis for specific inhibition of the highly sensitive ShHTL7 receptor. *Embo Rep* **19**, (2018).

50.	A. Hungler, A. Momin, K. Diederichs, S. T. Arold, ContaMiner and ContaBase: a webserver and database for early identification of unwantedly crystallized protein contaminants. *J Appl Crystallogr* **49**, 2252-2258 (2016).

51.	F. Long, A. A. Vagin, P. Young, G. N. Murshudov, BALBES: a molecular-replacement pipeline. *Acta Crystallogr D* **64**, 125-132 (2008).

52.	P. Emsley, K. Cowtan, Coot: model-building tools for molecular graphics. *Acta Crystallogr D* **60**, 2126-2132 (2004).

53.	P. V. Afonine *et al.*, Towards automated crystallographic structure refinement with phenix.refine. *Acta Crystallogr D* **68**, 352-367 (2012).

54.	J. Mi *et al.*, A manipulation of carotenoid metabolism influence biomass partitioning and fitness in tomato. *Metab Eng* **70**, 166-180 (2022).

55.	M. Jamil *et al.*, Striga hermonthica Suicidal Germination Activity of Potent Strigolactone Analogs: Evaluation from Laboratory Bioassays to Field Trials. *Plants (Basel)* **11**, (2022).

56.	J. Braguy *et al.*, SeedQuant: a deep learning-based tool for assessing stimulant and inhibitor activity on root parasitic seeds. *Plant Physiology* **186**, 1632-1644 (2021).
